# Anticipating critical transitions in epithelial-hybrid-mesenchymal cell-fate determination

**DOI:** 10.1101/733006

**Authors:** Sukanta Sarkar, Sudipta Kumar Sinha, Herbert Levine, Mohit Kumar Jolly, Partha Sharathi Dutta

## Abstract

In the vicinity of a tipping point, critical transitions occur when small changes in an input condition causes sudden, large and often irreversible changes in the state of a system. Many natural systems ranging from ecosystems to molecular biosystems are known to exhibit critical transitions in their response to stochastic perturbations. In diseases, an early prediction of upcoming critical transitions from a healthy to a disease state by using early warning signals is of prime interest due to potential application in forecasting disease onset. Here, we analyze cell-fate transitions between different phenotypes (epithelial, hybrid epithelial/mesenchymal (E/M) and mesenchymal states) that are implicated in cancer metastasis and chemoresistance. These transitions are mediated by a mutually inhibitory feedback loop microRNA-200/ZEB driven by the levels of transcription factor SNAIL. We find that the proximity to tipping points enabling these transitions among different phenotypes can be captured by critical slowing down based early warning signals, calculated from the trajectory of ZEB mRNA level. Further, the basin stability analysis reveals the unexpectedly large basin of attraction for a hybrid E/M phenotype. Finally, we identified mechanisms that can potentially elude the transition to a hybrid E/M phenotype. Overall, our results unravel the early warning signals that can be used to anticipate upcoming epithelial-hybrid-mesenchymal transitions. With the emerging evidence about the hybrid E/M phenotype being a key driver of metastasis, drug resistance, and tumor relapse; our results suggest ways to potentially evade these transitions, reducing the fitness of cancer cells and restricting tumor aggressiveness.

**Significance Statement:** Epithelial-hybrid-mesenchymal transitions play critical roles in cancer metastasis, drug resistance, and tumor relapse. Recent studies have proposed that cells in a hybrid epithelial/mesenchymal phenotype may be more aggressive than those on either end of the spectrum. However, no biomarker to predict upcoming transitions has been identified. Here, we show that critical slowing down based early warning signals can detect sudden transitions among epithelial, hybrid E/M, and mesenchymal phenotypes. Importantly, our results highlight how stable a hybrid E/M phenotype can be, and how can a transition to this state be avoided. Thus, our study provides valuable insights into restricting cellular plasticity en route metastasis.

## Introduction

Biological systems often display nonlinear dynamics and emergent complex behavior, and consequent multi-stability [1, 2]. This nonlinear behavior in many cases leads to ‘tipping points’ - threshold values at which the system abruptly shifts from one state to another, in response to small stochastic perturbations [3]. Such changes - referred to as critical transitions - have been observed in multiple instances of ecosystems, climate, financial markets [4, 5, 6], and more recently in many cases of health and disease [7, 1]. The consequences of critical transitions are often large and undesirable, for instance, the switch from a healthy state to a diseased state such as the onset of type-2 diabetes [8] or that of depression [9]. Moreover, these transitions are often difficult to reverse, potentially due to self-reinforcing positive feedback [10], thus, predicting the ‘tipping points’ can be crucial for preventing such catastrophic changes.

A critical transition is usually identified after a tipping point and is difficult to predict beforehand, because the equilibrium state of the system stays relatively unchanged until the tipping point is reached [1]. Thus, static observations may not be sufficient to predict these abrupt transitions. Many indicators of changing system dynamics have been suggested as early warning signals (EWS) for the impending critical transitions and have been experimentally shown to predict transitions in alternative states in yeast cultures [11] and plankton chemostats [12]. The most important clues for EWS arise from critical slowing down of the system as it approaches the tipping point. At the onset of a tipping point, the rate of return of the system to the current equilibrium state upon a random disturbance decreases as the dominant eigenvalue approaches zero, and eventually, this equilibrium state is replaced by the alternative state. Thus, under conditions of critical slowing down, the state of the system at a given time becomes increasingly like that at a previous moment, leading to higher temporal autocorrelation. Similarly, due to moving into a shallower well closer to the bifurcation point, the variance in data is increased [6]. Hence, two canonical statistical measures that are mostly used as EWS to indicate the proximity of a system to a tipping point are increasing variance and temporal lag-1 autocorrelation - AR(1) [3]. Few other measures used as EWS are recovery rate/return time [13, 12], skewness [14], conditional heteroskedasticity [15], spectral reddening [16], likelihood ratio [17] and interaction network based indicators [18].

While EWS and critical transitions have been well-studied in ecological and climate systems, their application in predicting disease onset is relatively recent and remains largely conceptual [1, 10]. Particularly, in cancer, critical transitions have been predicted in metabolic reprogramming [19] - a hallmark of cancer [20]. Here, we investigate critical transitions and EWS in another hallmark of cancer - invasion and metastasis. Metastasis - the spread of cancer cells from one organ to another - accounts for nearly all cancer related deaths in solid tumors [21]. Despite extensive genomic efforts, no specific mutational signatures have been yet identified for metastasis [22], thus limiting the druggable targets to restrict metastasis. Therefore, identifying tipping points for predicting and preventing metastasis can be beneficial in curbing tumor aggressiveness.

Most solid tumors originate in epithelial organs where cells do not typically migrate or invade, rather maintain tight cell-cell adhesion and a specific tissue organization. Thus, to metastasize, they typically undergo a phenotypic switch known as epithelial-mesenchymal transition (EMT) where they lose cell-cell adhesion and gain the traits of migration and invasion [23]. Cells undergoing EMT get launched into the bloodstream, and also gain the ability to initiate new tumors at metastatic sites, gain resistance against multiple drugs [24], and evade attacks by the immune system [25]. Thus, EMT provides multiple survival advantages to disseminated cells that typically undergo a mesenchymal-epithelial transition (MET) to colonize distant organs. Recent investigations, including ours, have identified that EMT and MET need not be binary processes, instead cells can undergo partial EMT/MET and stably maintain one or more hybrid-epithelial/mesenchymal (E/M) phenotype(s) [23]. Importantly, cells in hybrid-E/M phenotype(s), i.e. those that undergo partial EMT, may be even more aggressive than cells that have undergone full EMT [26, 27]. However, no specific biomarker has been identified that can a priori predict the onset of transitions among epithelial, mesenchymal and hybrid-E/M states. Thus, identifying EWS for transitions among these cell states can be a valuable contribution towards restricting them.

Here, we identify critical slowing down based EWS in a core regulatory network of EMT/MET. Three well known indicators - lag-1 autocorrelation, variance and conditional heteroskedasticity - work well to forewarn upcoming transitions among epithelial, hybrid-epithelial/mesenchymal, and mesenchymal states, thus opening the possibility of considering EWS as biomarkers to forewarn cancer metastasis. We also calculate the basin stability measure to evaluate the probability of occurrence of a particular state in various multistable regions. A higher basin stability measure corresponding to a particular state determines larger possibility of attaining the state in a multistable region. Complementing our basin stability measures with potential landscapes and phase diagrams for EMT circuit, we identify how a monostable hybrid E/M state can be maintained and thus suggest mechanisms to avoid it. Overall, our results highlight the ability to predict cellular transitions in metastasis before they occur and may provide a dynamic biomarker to gauge metastatic potential.

## Model

We consider an analytical model of microRNA (miR) based chimeric circuit developed by Lu et al. [28]. The model incorporates the features of miR mediated regulation in the translation-transcription processes and captures the formation of various miR-mRNA complexes by the binding/unbinding dynamics of miR and mRNA (see Fig. 1A). The deterministic equations of the circuit which govern the combined dynamics of miR (*μ*), mRNA (*m*) and TF protein (*B*) are given by:

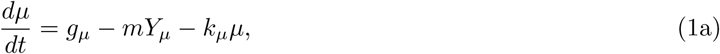

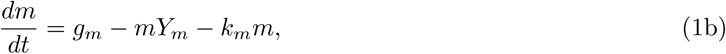

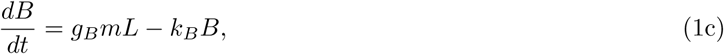

where *g*_*μ*_ and *g*_*m*_ are the synthesis rates of *μ* and *m*, respectively, and *g*_*B*_ is the translation rate of protein *B* for each *m* in the absence of *μ. k*_*μ*_, *k*_*m*_ and *k*_*B*_ are the degradation rates of *μ, m* and *B*, respectively. *Y*_*μ*_, *Y*_*m*_ and *L* are *μ* dependent functions [28] denoting various effects of microRNA-mediated repression.

**Figure 1.**
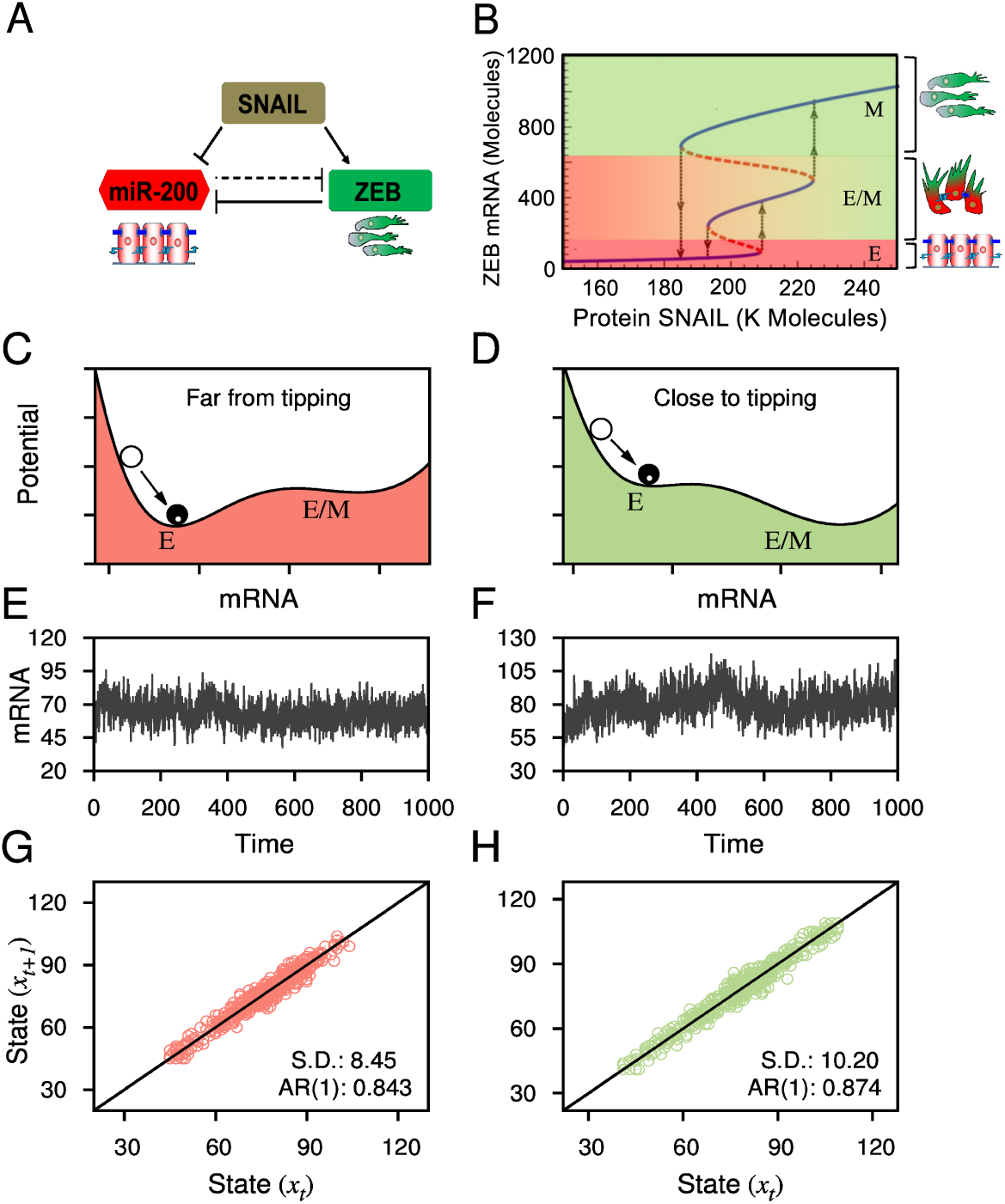
(A) Schematic diagram of the microRNA-based chimeric circuit. (B) Bifurcation diagram depicting the changes in ZEB mRNA levels with variations in the levels of SNAIL. E, hybrid E/M and M denote epithelial state, hybrid epithelial/mesenchymal state and mesenchymal state, respectively: lowest levels of ZEB mRNA correspond to epithelial state, intermediate levels to a hybrid E/M state, and highest ones to mesenchymal state, as shown by corresponding cartoons. (C-F) An overview of critical transition in the circuit which has multistability. Schematic potential landscapes representing two stable states (i.e. E and hybrid E/M) of deterministic system: (C) high resilience of the E state when it is far from the tipping point, and (D) low resilience of the state close to a tipping point, when the system approaches a sudden shift from E to hybrid E/M state. Stochastic time series of the system (2): (E) with S=197K (far from the tipping point) and (F) with S=207K (close to the tipping point), respectively. (G, H) In the vicinity of a tipping point, due to decreasing resilience the system has stronger memory for perturbation in comparison to that of far from a tipping point and that are characterized by larger standard deviation (S.D.) and lag-1 autocorrelation (AR(1)). All other parameters for the circuit are given in SI.

The corresponding chimeric tristable miR-200/ZEB circuit is modeled as:

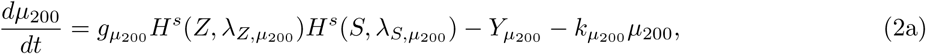

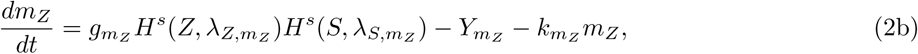

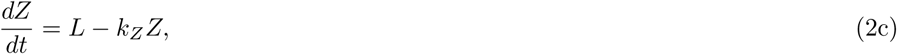

where *H*^*s*^ is the Hill function (details are in *SI Text, Sections 1 and 2*).

As a stochastic description of Eqs. (2) can accurately capture the dynamics of the system, we derive the corresponding chemical Master equation which follows from birth-death processes [29]. The Master equation is given by:

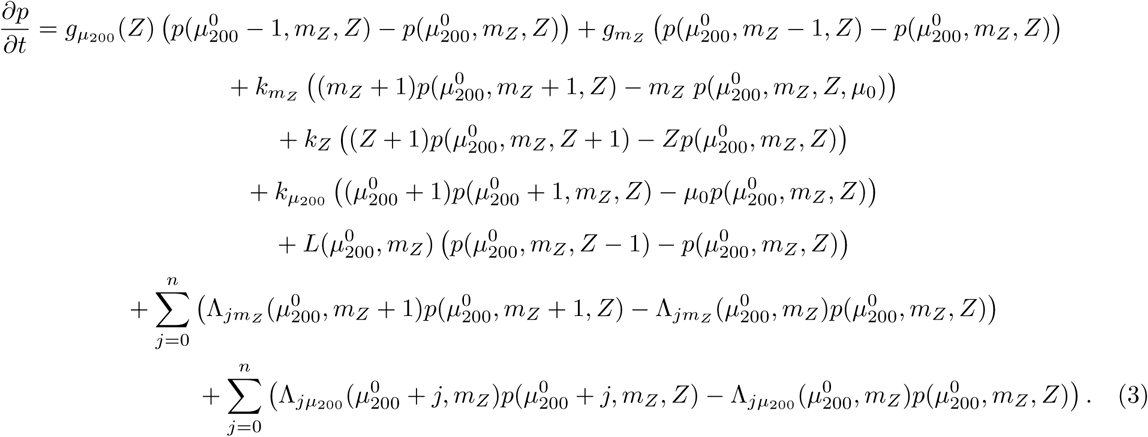

Where 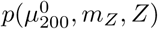 is the grand probability function. The Eq. (3) is a birth-death process for the probabilities of the separate states specified by the values of 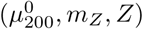. All the terms appear in the equation as pairs: (i) birth of a state 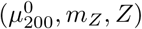 due to transition from other states 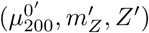, and (ii) death due transition from 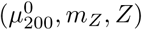 into other states. There are ten such processes associated with birth and death of miR, mRNA and ZEB in our model (see also *SI Text, Section 3* for details). We have simulated this Master equation with Gillespie algorithm [30] to obtain the stochastic trajectory of the system (*SI Text, Section 3A*). The stochastic trajectory of the system identifies the occurrence of critical transition between different phenotypes and using critical slowing down based EWS we are able to forecast such transitions beforehand.

## Results and Discussion

### Bifurcation-induced tipping signs in epithelial-hybrid-mesenchymal transition

A mutually inhibitory feedback loop between members of ZEB transcription factor and those of microRNA (miR)200 has been postulated to govern EMT/MET; ZEB can drive EMT by inhibiting cell-cell adhesion and cell polarity, while miR-200 tend to maintain an epithelial phenotype [23]. Unlike mutually inhibiting feedback loops where both players are transcription factors, this loop is chimeric, i.e. it contains both transcriptional and translational regulation [28, 31]. First, we perform the bifurcation analysis of this deterministic tristable chimeric circuit Eqs. (2) with variations in the SNAIL concentration (S) (see Fig. 1B). The values of all the other model parameters of this circuit are presented in the *SI Text, Table S1-S3*. We denote three coexisting stable states: (high miR-200/low ZEB), (low miR-200/high ZEB), and (medium miR-200/medium ZEB). These states correspond to epithelial (E) and mesenchymal (M), and hybrid-epithelial/mesenchymal (E/M) phenotypes respectively [32, 23]. For increasing levels of S, the circuit first exhibits monostable E state; an increase in S leads to bistability between E and M states; a further increase enables tristability between E, hybrid-E/M and M states; then bistability between hybrid-E/M and M states, and finally a monostable M state. The existence of multistable regions includes the appearance of saddle-node bifurcations and hysteresis loops that triggers the possibility of occurrence of catastrophic critical transitions in the presence of intrinsic stochastic perturbations [33].

Since this feedback loop exhibits tristability, it may pass through two critical points (or tipping points) and, therefore can reach two alternative states, one after another. A systematic analysis of such critical transition is commonly done by analysing stochastic trajectory. In Fig. 1, we have presented a brief overview of critical transition in the EMT circuit from pure E to hybrid-E/M phenotype transition with variations in the levels of protein SNAIL, when the system is far from or close to a tipping point (see Figs. 1C,D). More specifically, larger variance and increased lag-1 autocorrelation determine the proximity to a tipping point (see Figs. 1G,H). With increasing SNAIL value the system may experience two subsequent transitions, one from E to hybrid-E/M state and another from hybrid-E/M to M state. However, while decreasing SNAIL value results in a direct transition from M to E state which bypasses the hybrid-E/M state.

### Early warning signals for transitions among epithelial, hybrid-E/M and mesenchymal states

We began our search for signals of critical slowing down by calculating EWS of critical transitions in data sets obtained from stochastic simulations (see *Materials and Methods section*) of the chimeric circuit. The stochastic trajectory (time series) representing ZEB mRNA levels, with continuously increasing SNAIL value, exhibits sudden transitions from E state to hybrid-E/M state and further hybrid-E/M state to M state (See Fig. 2A). The trajectory is generated with time varying signal SNAIL. The SNAIL level starts at 150K molecules at day 0 and then increases upto 250K molecules at day 20. This increase in SNAIL levels can drive EMT in a cell, i.e. moving from monostable epithelial region to a monostable mesenchymal region (Fig. 1B), and the timescale over which SNAIL levels are varied are commensurate with those over which EMT is observed [34, 35].

**Figure 2.**
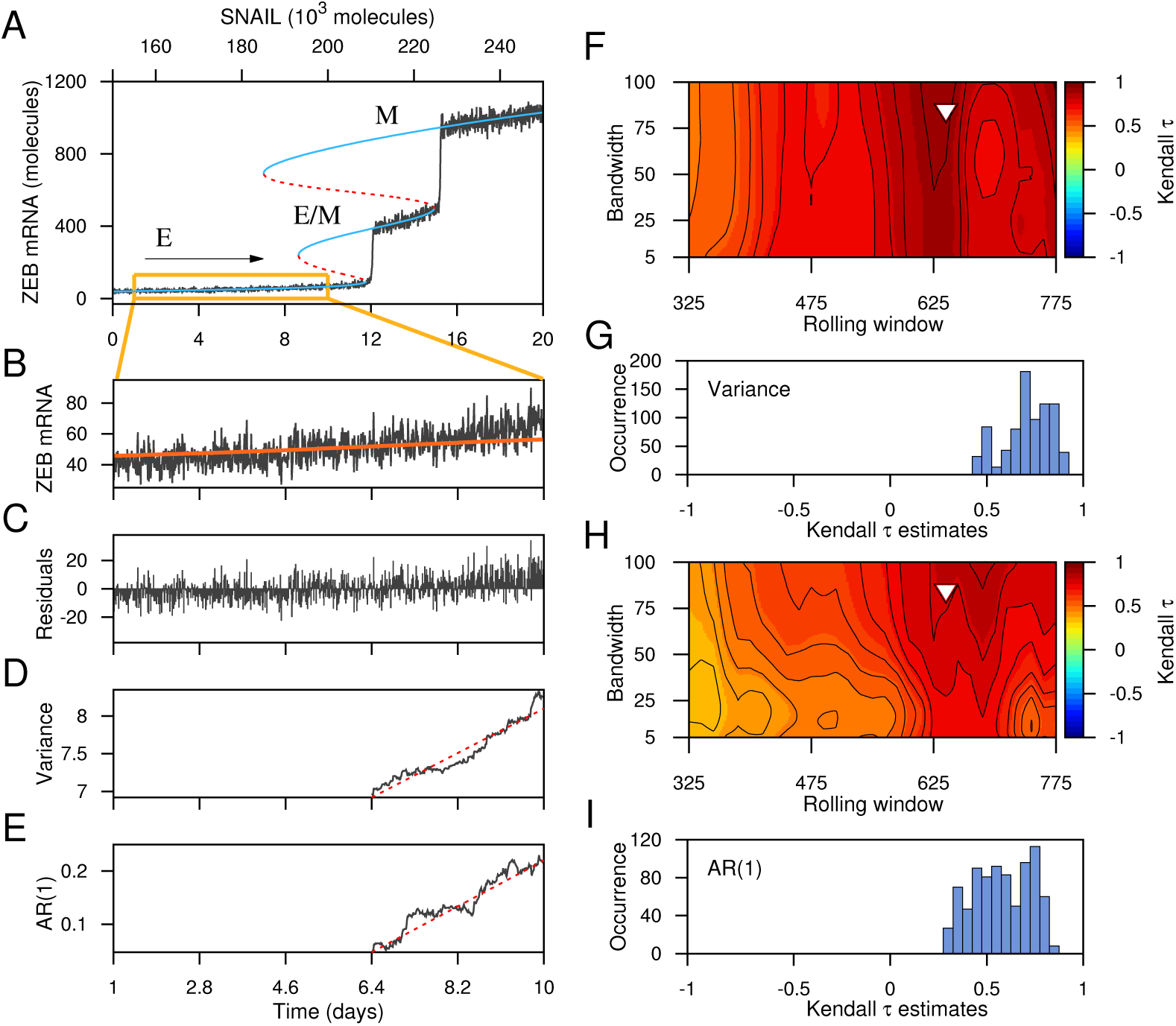
Critical transitions between different cell states of the regulatory circuit that are driven by *forward* change in the control parameter SNAIL, and indicators of critical slowing down. (A) Transitions from E state to hybrid E/M state and hybrid E/M state to M state. (B) Stochastic time series segment of the system before the transition to hybrid E/M state (a segment as indicated by the boxed region in (A)). (C) Residual time series after applying Gaussian filter (red curve in (B) is the trend used for filtering). EWS calculated from the filtered time series after using a rolling window of 60% of the data length: (D) variance and (E) AR(1). (F-I) Sensitivity analysis of the filtering bandwidth and the rolling window size used to calculate the EWS. Contour plots reveal the effect of variable rolling window size and filtering bandwidth on the observed trend in the EWS, (F) variance and (H) AR(1), for the filtered data as measured by the Kendall-*τ* value. The triangles indicate the rolling window size and bandwidth used to calculate the EWS. Frequency distributions of Kendall-*τ* values for (G) variance and (I) AR(1).

First we evaluate the effectiveness of different EWS to positively alarm an impending sudden catastrophic transition from E state to hybrid-E/M state, by tracking the values of ZEB mRNA. For EWS analysis, we consider a time series segment before the transition to hybrid-E/M state (see Fig. 2B). To filter possible non-stationarities in the data we subtracted a Gaussian kernel smoothing function across the time series segment and used the remaining residuals (Fig. 2C) for EWS analysis [36]. We calculate the variance and lag-1 autocorrelation (AR(1)) (see *SI Text, Section 4*) values with a rolling window having a length of 60% the length of the residual time series segment and found both the variance and AR(1) value to be increasing (see Figs. 2D-E). A concurrent increase in the EWS is an well known indicator of an upcoming critical transition [3, 5]. The performance of EWS is in general known to be sensitive to the choice of the filtering bandwidth used in Gaussian kernel smoothing and also on the rolling window size [37, 38]. The bandwidth of kernel smoothing determines the degree of data smoothing without filtering the low frequencies from the data and the choice of rolling window size depends on a trade-off between data resolution and reliability of the estimation of EWS. Therefore, rather than choosing arbitrary values, here we perform sensitivity analysis, of the filtering bandwidth and rolling window size (see Figs. 2F-I). For sensitivity analysis the rolling window size was varied from 25% to 75% of the data length in increments of 15 points, together with variations in the filtering bandwidth ranging from 5 to 100 in increments of 10. For all possible combinations of these two parameters, the observed trends in variance and AR(1) were quantified using the non-parametric Kendall’s *τ* rank correlation coefficient. A positive Kendall’s *τ* determines increasing trend in the EWS prior to a critical transition. To maximise the estimated trends for the EWS, we have used the sensitivity plot to select a particular filtering bandwidth and window size (see Fig. 2F for variance and Fig. 2H for AR(1)) (for details see *SI Text, Section 4B*). The frequency distributions of the Kendall’s trend statistic for the variance and the AR(1) are presented in Fig. 2G, I, respectively.

The EWS work well for capturing the transition from hybrid-E/M state to M state (see *SI Text, Section 5*), suggesting that transitions in the forward direction (i.e. increase in SNAIL) can be captured by stochastic time series of ZEB mRNA. We generate the stochastic time series of ZEB mRNA from the probabilistic model through the Monte Carlo simulations [30] which incorporates intrinsic cellular noise. We vary both the time and the parameter (the number of SNAIL molecules) together, which carries the signature of critical slowing down while shifting to an alternative stable state. We carried out our simulations for a period of 0 to 20 days along with the simultaneous variations in the number of SNAIL molecules, that varies from 150K to 250K molecules.

Next, we investigated whether these EWS can also be observed in backward transitions, i.e. with decreasing value of SNAIL (Fig. 3). Due to the hysteresis and asymmetry in transitions in both directions (E to M vs. M to E), we observe sudden direct transition from M to E state (see Fig. 3A) bypassing the hybrid-E/M state. We consider a time series segment prior the transition to E state (Fig. 3B) and further used the residual time series for EWS analysis (Fig. 3C). Importantly, both the EWS markers - variance and AR(1) - shown an increasing trend closer to the tipping point for this transition from M to E (see Figs. 3D-E). Reinforcing our previous analysis, these EWS were evaluated with specific choices of detrending bandwidth and rolling window size to maximise their trends. Put together, these results highlight that the transitions among E, hybrid-E/M and M states can be predicted before they occur, using EWS variance and AR(1).

**Figure 3.**
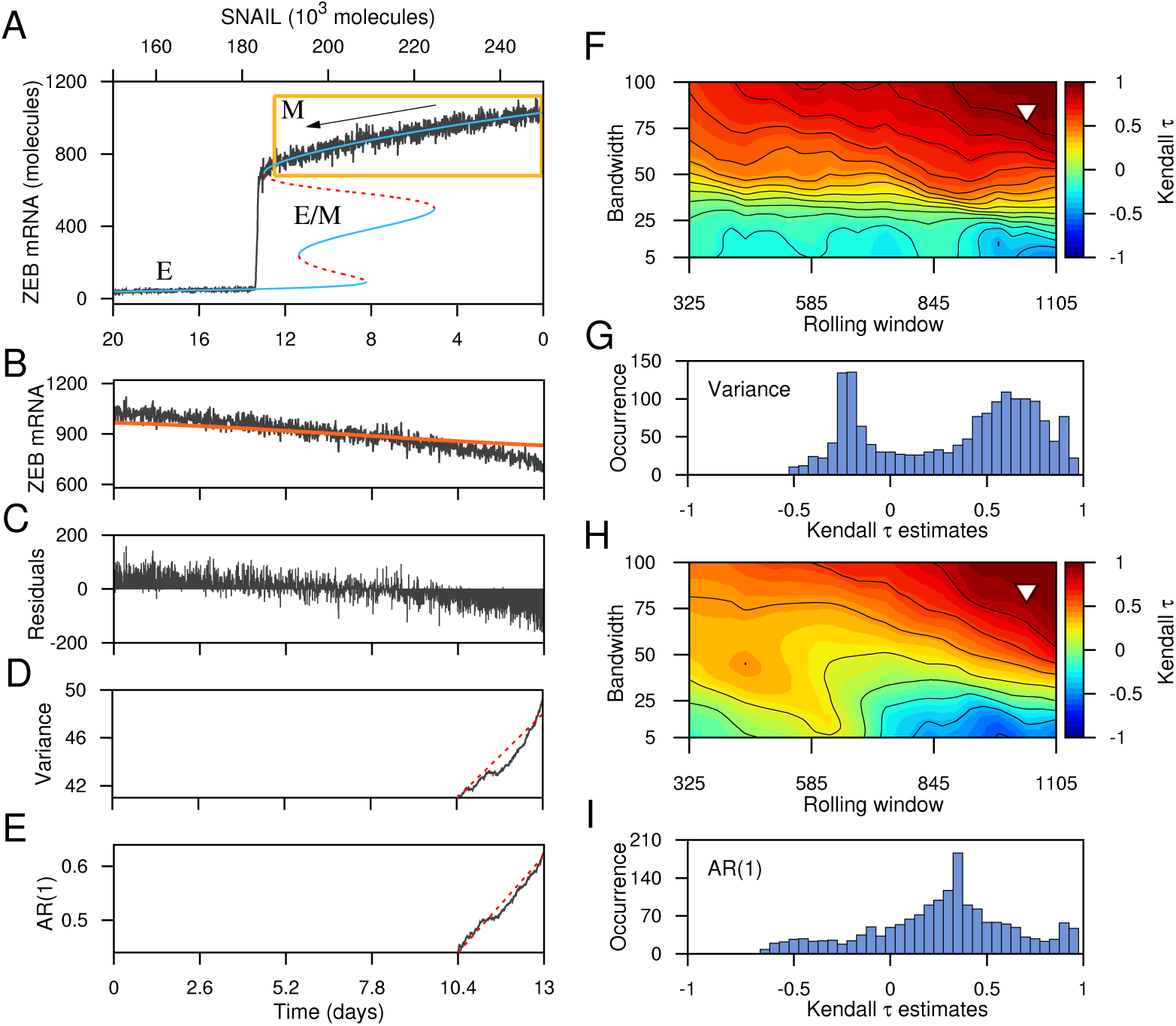
Critical transition between different cell states of the regulatory circuit that is driven by *backward* change in the control parameter SNAIL, and indicators of critical slowing down. (A) Transition from M state to E state that bypasses the hybrid E/M state. (B) Stochastic time series segment of the system before the transition to E state (a segment as indicated by the boxed region in (A)). (C) Residual time series after applying Gaussian filter (red curve in (B) is the trend used for filtering). EWS calculated from the filtered time series after using a rolling window of 80% of the data length: (D) variance and (E) AR(1). Contour plots reveal the effect of variable rolling window size and filtering bandwidth on the observed trend in the EWS, (F) variance and (H) AR(1), for the filtered data as measured by the Kendall-*τ* value. The triangles indicate the rolling window size and bandwidth used to calculate the EWS. Frequency distributions of Kendall-*τ* values for (G) variance and (I) AR(1).

Further, for the aforementioned three transitions, E to hybrid E/M, hybrid E/M to M and M to E state, we evaluate the robustness of EWS trends to all the rolling window sizes depicted as the distribution of the Kendall-*τ* statistic around their median, for both the ‘original’ and ‘surrogate’ time series (see *SI Text, Section 6* and *SI Fig. S2*). In the case of ‘original’ data sets, most of the trends for AR(1) and variance are robust to rolling window sizes as majority of the associated box-plots stays above the *y*-zero axes [39].

### Conditional heteroskedasticity applied as early warning signals

To evaluate robustness of the predictions made by the EWS variance and AR(1), we calculate conditional heteroskidasticity - one of the other measures known to forewarn critical transitions [15]. Conditional heteroskidasticity is indicated by the persistence in the conditional variance of the error term in time series models [40]. The advantage of this indicator over others is that it minimizes the chance of the occurrence of false positive signals in time series that does not have any critical transition. To calculate conditional heteroskidasticity, time series is modelled as an auto-regressive process and the residuals are obtained. The persistence of the conditional variance of the residuals then determine the conditional heteroskidasticity (see *SI Text, Section 4C* for details of the procedure). Prior to a critical transition, significant conditional heteroskidasticity is expected to be visible in the time series [15].

We consider the time series segments before the critical transitions for both the cases; E to hybrid-E/M transition and M to E transition (see Fig. 2B and Fig. 3B). Figure 4A presents the squared residuals from an auto-regressive lag-1 model applied to the time series segment of E to hybrid-E/M transition (Fig. 2B) plotted with the residuals at the next time step. The slanted line is the regression line. The positive correlation between the squared residuals at time step *t* and time step *t* + 1 indicates conditional heteroskidasticity. We also apply the cumulative number of significant Lagrange multiplier test (*C*) to the time series (Fig. 4B). The cumulative increases prior to the critical transition indicating that significant number of tests shows conditional heteroskedasticity in the time series. For the transition to M to E state, we get similar result (Fig. 4C, D).

**Figure 4.**
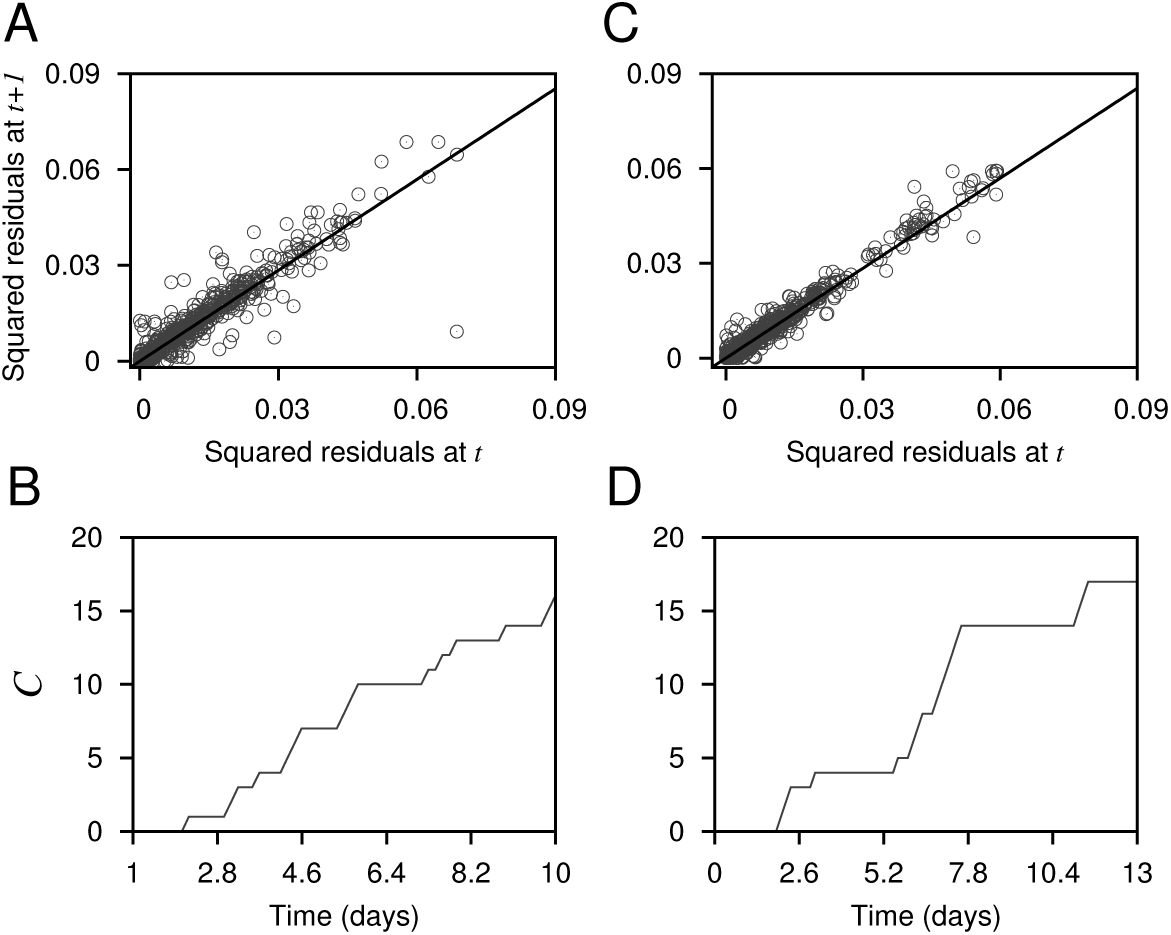
(A, C) Squared residuals from an autoregressive lag-1 model plotted with the next squared residuals and (B, D) cumulative number of significant Lagrange multiplier test (*C*), both applied to the data presented in Fig. 2B (for A, B) and Fig. 3B (for C, D), respectively. In (A, B), the black slanted lines are fitted regression lines at lag-1.

### Stochastic potential and basin stability analyses reveal relative stability of the three cell states

For a dynamical system, a potential well represents the existence of a steady state. Here, we projected the stochastic potential of the system in ZEB mRNA – *μ*RNA200 plane for different values of the parameter SNAIL (Fig. 5). The lowest value of the potential corresponds to the existence of a deep well and hence subsequently the existence of a steady state. Here, for different SNAIL values, the stochastic potentials clearly exhibit bistable/multistable states. Consistent with the deterministic dynamics of the system (Fig. 1B), we note the co-existence of E and M states (Fig. 5A), the co-existence of all the three E, hybrid-E/M and M states (Fig. 5B), and the co-existence of hybrid-E/M and M states (Fig. 5C). The details of the method used to calculate the stochastic potentials are given in the *SI Text, Section 7*.

**Figure 5.**
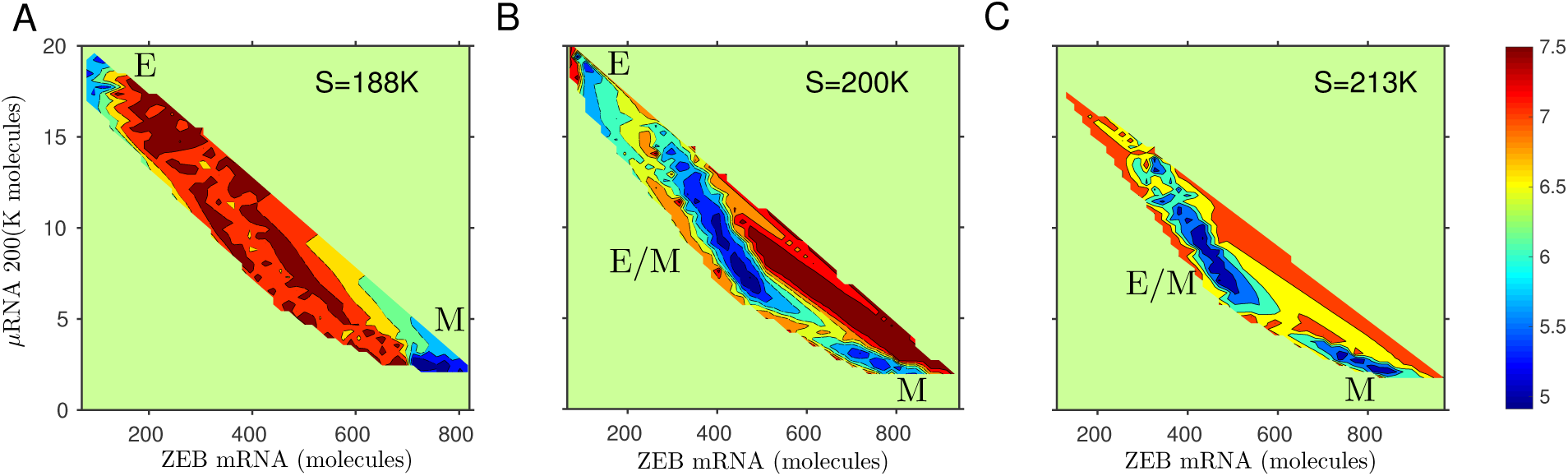
The potential landscapes of the genetic circuit in two-dimensional mRNA-*μ*RNA plane for different values of SNAIL (S). The blue regions represent lower potential and correspondingly higher probability of occurrence of a steady state. Identifying the existence of: (A) epithelial and mesenchymal states at S=188K, (B) epithelial, hybrid epithelial-mesenchymal and mesenchymal states at S=200K, and (C) hybrid epithelial-mesenchymal and mesenchymal states at S=213K.

Given that the hybrid-E/M state has been proposed to be the ‘fittest’ for metastasis [41] and that we observed a relatively larger region denoting the stability of hybrid-E/M in the tristable region (Fig. 5), we investigated the probability of attaining the hybrid-E/M state in a tristable region in the presence of random perturbations. This probability can be calculated by performing basin stability measure [42].

For a complex system, basin stability is a measurement of the stability/resilience of a steady state in a probability sense which pivots on the volume of the basin of attraction. In other words, it measures the likelihood of return to a steady state after random, non-small perturbations. Thus, for a high-dimensional multistable system, it is a powerful tool to measure the basin volume (see *SI Text, Section 8*). For our system, we observe multistability for different parameter values of SNAIL(S). For S=188K (see Fig. 1B), the system has coexisting E and M states. Basin stability measures that for a sufficiently large set of random initial conditions, E and M states have probabilities 0.91 and 0.09 of return to their original state, i.e. among all random initial conditions 91% and 9% trajectories will reach E and M states (Fig. 6A), respectively. For S=200K, system have probabilities 0.1, 0.76 and 0.23 of reaching to E, hybrid-E/M and M states (Fig. 6B), respectively from a set of random initial states. Similarly for S=213K, the corresponding probabilities of return to hybrid-E/M state and M state are 0.62 and 0.38 (Fig. 6C), respectively. Further, increase in the levels of SNAIL at S=220K reduces the probability of attaining hybrid-E/M state which becomes 0.52 and remaining 0.48 is the probability of attaining the M state (Fig. 6D), indicating that as we proceed from a bistable M-E/M phase to a monostable M phase, the basin stability of E/M decreases, being conceptually consistent with the mean residence time analysis for this circuit [43].

**Figure 6.**
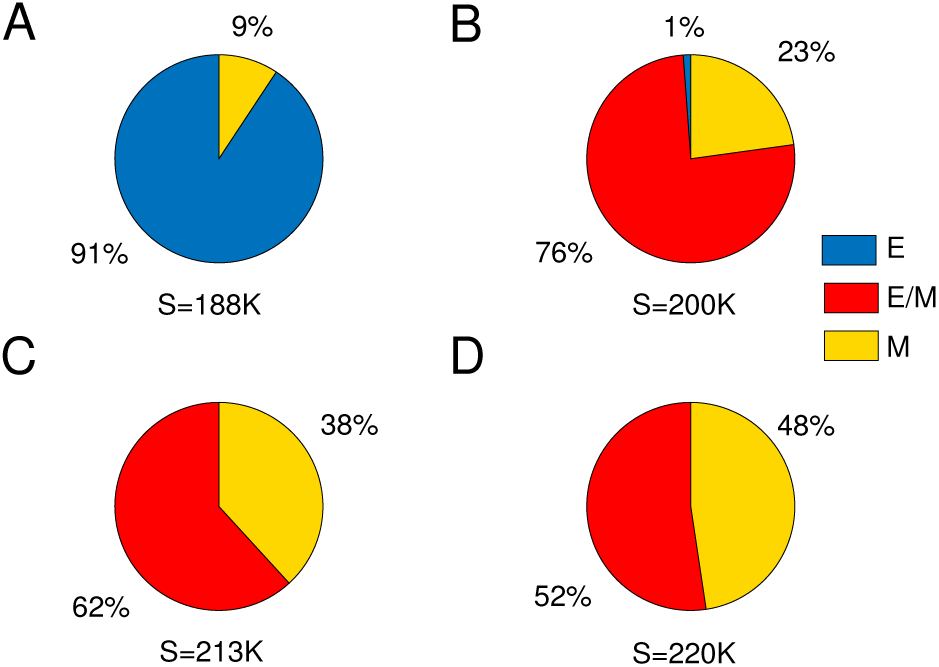
Pie diagrams representing basin stability of the system for different values of SNAIL (S): (A) S=188K, (B) S=200K, (C) S=213K and (D) S=220K. The percentage of 10^4^ simulations with random initial conditions reaching to a particular steady state in a bistable/multistable region. Blue, red and yellow regions correspond to the % of simulations reaching to any one of the E, hybrid-E*/*M and M states, respectively.

Hence, the basin stability results suggest that an E state is more stable in bistable region containing both E and M states, but the hybrid E/M state is more stable for the two later cases. Thus, in the (miR-200/ZEB) loop, chances of getting a hybrid E/M state seems relatively very high compared to the other two states (see *SI Fig. S3*). This result is reminiscent of mean residence times calculations for E, hybrid-E/M and M states [43], and suggests that hybrid-E/M state is not perhaps as ‘metastable’ as was initially postulated experimentally [23].

### Identifying mechanisms to evade the transition into aggressive hybrid E/M state

Next, we sought after mechanisms to evade transition to a hybrid-E/M state, given its association with higher aggressiveness and worse patient survival. We first identified what mechanisms can lead to stabilized hybrid-E/M state. So far, our results have identified monostable E, monostable M, and other bistable and tristable regions, but not a monostable hybrid-E/M state. Including other factors such as GRHL2, NUMB in the network can enable the existence of a monostable hybrid-E/M region [27]. Here, we analyzed the parameter space of the miR-200/ZEB feedback loop to identify regions enabling the existence of a hybrid-E/M state as a monostable phase, without adding more components in the network. We varied the levels of SNAIL, and the threshold (half-maximal concentration) value of ZEB in the shifted Hill function corresponding to ZEB inhibiting miR-200, and calculated the phase diagram shown in Fig. 7. The different phases in the diagram are separated by four saddle-node bifurcation curves. We could identify a large parameter region in which the monostable hybrid-E/M phase appears - high levels of both SNAIL and the threshold of ZEB (see Fig. 7A). This result suggests that as the strength of inhibition of miR-200 by ZEB is weakened, the progression to a complete EMT may be halted and cells can stably occupy a hybrid-E/M state for higher values of SNAIL (Fig. 7A). Conversely, as this inhibition is made stronger, the stability of the hybrid-E/M state gradually decreases (Fig. 7C) and eventually the hybrid-E/M state disappears (Fig. 7B). Here, the hybrid-E/M state disappears when the systems response curve changes from ‘folded’ to ‘smooth’, in response to the variations in the input condition. In fact the folded response curve looks like a typical first-order (i.e. abrupt) or discontinuous transition (Fig. 7C), however contains two unstable states and one stable hybrid-E/M state which in general shows two stable and one unstable states in most of the studies on critical transitions [44, 45]. The smooth response curve corresponds to second-order (i.e. continuous) phase transition that has only one unstable state here (Fig. 7B), in contrast to a bistable system which has a stable state. Therefore the dynamical mechanism behind the disappearance of the hybrid-E/M state is the changeover from first-to second-order phase transition in the systems response curve.

**Figure 7.**
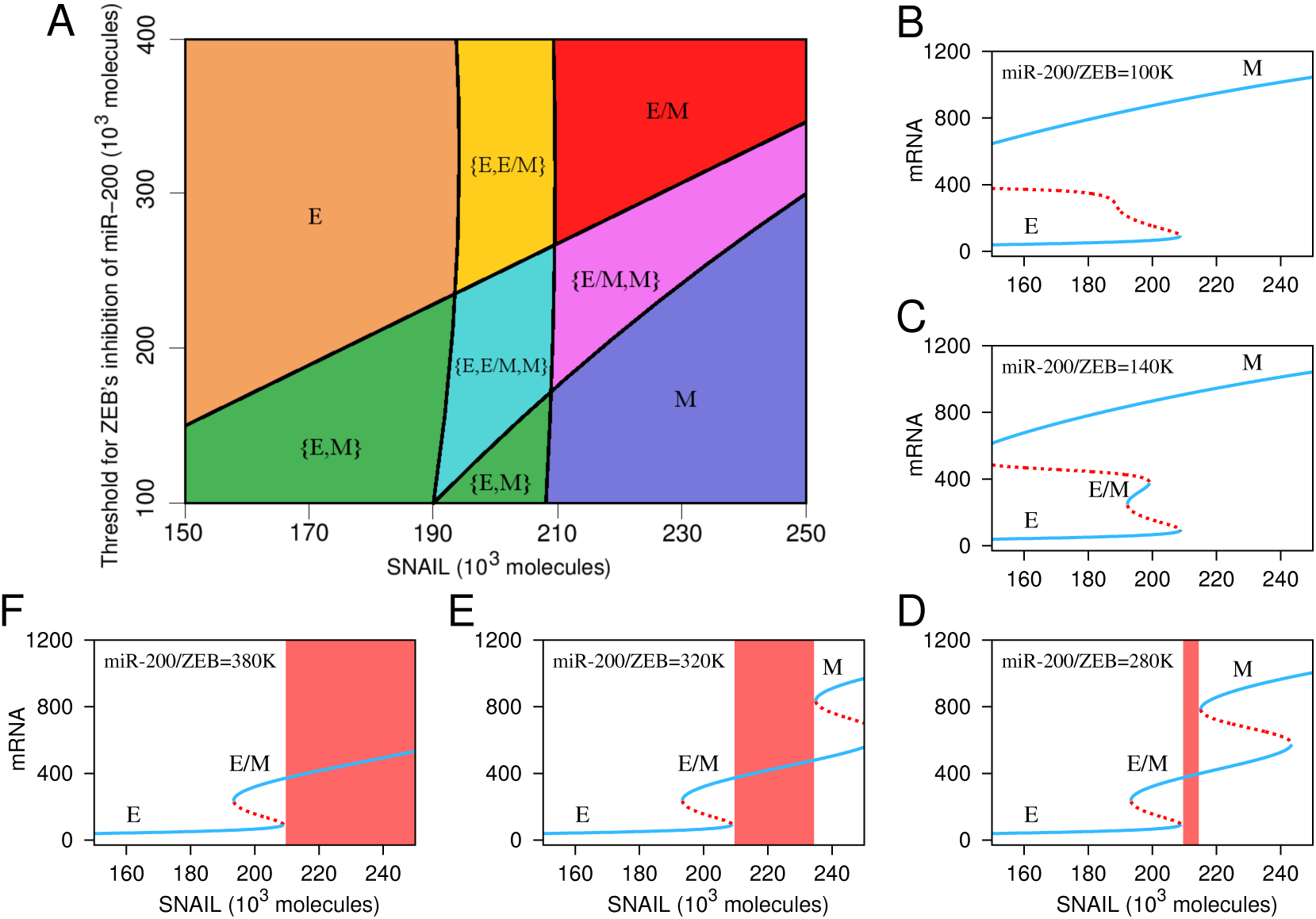
The phase diagram and corresponding bifurcation diagrams of the genetic circuit. (A) The phase diagram of the genetic circuit with variations in SNAIL and miR-200/ZEB levels. Each phase corresponds to either any of the monostable state or coexisting bistable/multistable states. For example, in E the epithelial state is stable, in {E, hybrid-E/M} both epithelial and hybrid-epithelial mesenchymal states coexist. (B-F) Bifurcation diagrams of mRNA with variations in the level of SNAIL for different values of miR-200/ZEB: (B) 100K, (C) 140K, (D) 280K, (E) 320K and (F) 380K. As we move from (D) to (F), the monostable region for hybrid-E*/*M state increases.

Similarly, we also varied the levels of SNAIL and the threshold of self-activation of ZEB (see *SI Text, Section 9*). Reduced threshold, i.e. stronger self-activation, enable a monostable hybrid E/M phase at lower SNAIL values, while it disappears with increased threshold, i.e. weaker self-activation of ZEB (see *SI Fig. S4*). Increasing SNAIL values and weakening self-activation drive the system towards a bistable E, M phase, i.e. disappearance of E/M state. These results suggest that a balance between strengths of mutual inhibition and self-activation can enable the existence of a hybrid E/M phenotype [31].

## Discussion

Anticipating critical transitions remains an extremely challenging task in multiple scenarios including eutropication of lakes, crash of financial markets, and more importantly, in onset of disease. The system typically displays almost no sign of the impending transition until it happens, thus using early warning signals (EWS) such as variance, autocorrelation and conditional heteroskedasticity can be used to forecast the critical transitions which are often catastrophic. Here, we show that these EWS can capture the transitions among epithelial, mesenchymal and hybrid epithelial/mesenchymal phenotypes. This phenotypic plasticity drives cancer metastasis and drug resistance in cancer - the cause of almost all cancer-related deaths. Given that no unique mutational signature has been yet identified for metastasis, despite extensive genomic efforts, these EWS that can predict the onset of these cellular transitions that govern metastasis can serve as potentially important dynamic biomarkers. Recent efforts have focused on identifying such dynamic biomarkers in the context of pulmonary metastasis of hepatocellular carcinoma [46]. With more single-cell dynamic data emerging in the context of epithelial-hybrid-mesenchymal transitions [47], using EWS signals can help predict the tipping point of metastasis initiation.

Cancer metastasis has been long thought to be driven solely by individual cell migration (i.e. a mesenchymal state), however, recent studies have questioned this dogma, highlighting that not only clustered cell migration can be possible during metastasis, but also it can be the predominant driver of metastasis [48, 49]. These clusters, typically 5-8 cells large, can pass through capillaries by arranging themselves transiently into a single-file chain [50], and can contain non-cancerous cells that can facilitate metastasis [51]. A hybrid-E/M phenotype has been associated with such collective/clustered cell migration [52, 53], thus, our analysis identifying the relatively high basin stability of the hybrid-E/M phenotype can help explain the ability of cancer cells to form clusters of circulating tumor cells.

Here, our analysis focused on temporal dynamics of a gene regulatory network for EMT; however, EWS have also been identified in spatiotemporal dynamics, particularly in ecology [54, 55]. Thus, EWS can also be potentially identified in a spatially extended regulatory networks for EMT, for instance, investigating the varying extents of EMT induction in different parts of a tissue [56, 57] or identifying critical transitions in cancer-immune interplay [58]. Further, besides EMT, there are other axes of phenotypic plasticity in cancer, such as metabolic plasticity, switching back and forth between a cancer stem cell (CSCs) and a non-cancer stem cell. With recent developments in identifying the multistable dynamics of the networks regulating these transitions [59]. EWS analysis can be applied to these networks to identifying promising novel dynamic biomarkers. However, an open question remains: can we identify the strongest and most robust signal of critical transition, among many which might show EWS? For instance, during metastasis, players involved in EMT, CSCs, and metabolic plasticity may all show EWS and vary dynamically, but which among these interconnected axes can be considered as the Achilles’ heel of metastatic potential needs to be identified rigorously?

Majority of the critical slowing down based EWS used to predict critical transitions in natural systems involves saddle-node bifurcation under the presence of white noise (temporally uncorrelated noise) that perturbs the abundance of the system [60]. For a large class of systems that exhibit other type of bifurcations apart form the saddle-node, the effectiveness of EWS remains largely unknown. For different type of bifurcations (e.g. transcritical, pitchfork, supercritical Hopf bifurcation) with diverse noise (e.g. coloured (temporally correlated) noise) EWS do not always work reliably to forecast sudden critical transitions [61]. They found to be very sensitive to the length of pre transition time series data, and also to other decisions like filtering bandwidth and rolling window size [38]. There also exist systems in which bifurcations occur without critical slowing down, such as in a structured consumer resource model where the upper point equilibrium coexists with a lower limit cycle [62], occurrence of basin boundary collisions [63] and as a result in these systems EWS do not work properly. In fact in general EWS work well for the situations when critical transition and critical slowing down co-occur [61]. Although robustness of EWS have been successfully shown in some cases [3, 60], a detail analysis of their effectiveness is still an open challenge [17, 64, 61].

In summary, our analysis strongly indicates the presence of EWS during epithelial-hybrid-mesenchymal transitions - a central motor of cellular plasticity during cancer metastasis and emergence of therapy resistance [65]. We show that many robust measures of EWS - increased variance, autocorrelation and conditional heteroskedascity - vary dynamically as cells transition among these three phenotypes. Our results also identify increased basin stability of a hybrid-E/M phenotype - considered to be the ‘fittest’ for metastasis - and suggests ways how to elude transitions into the hybrid-E/M state, potentially restricting cancer spread.

## Materials and Methods

### Numerical simulations and bifurcation diagrams of the deterministic system

We have used Matlab (R2015b) for numerical simulations of the deterministic system (Eq. 2). The codimension-one bifurcation diagrams involving two or more saddle-node bifurcation points were obtained using the continuation package MATCONT [66]. The two parameter bifurcation diagram (i.e. the phase diagram) with variations in the parameters SNAIL and miR-200/ZEB was obtained through the calculations of multiple codimension-one bifurcations points. Later, the bifurcation curves separating monostable, bistable and tristable existence regions of the steady states were presented by connecting multiple codimension-one bifurcations points.

### Stochastic system and Monte Carlo simulations

The time series of ZEB mRNA levels was generated from the probabilistic model through Monte Carlo simulations [30] which incorporates the intrinsic cellular noise. The algorithm considers each of the reaction events as individual realisations of Markov process. The time and species numbers are updated stochastically by choosing a random reaction event. The miR(*μ*) based chimeric tristable miR-200/ZEB circuit is simulated by realising ten reaction events as a function of the number of SNAIL molecules. The reaction events are listed in the *SI Table S4*. All biochemical parameters are based on [32] and those are listed in the *SI Table S1, S2 and S3* for completeness. Both the time and the parameter (number of SNAIL molecules) are varied together to obtain the time series of ZEB mRNA levels that carries the signature of critical slowing down while shifting to an alternative stable state. In particular, we perform our simulations for a period of 20 days along with the simultaneous variations in the number of SNAIL molecules, that ranges from 150K to 250K molecules. The chosen time period and and the range of SNAIL molecules are in consistent in the context of epithelial to mesenchymal transition period [32]. More details of the simulation is presented in the *SI Text, Section 3*.

### Statistical analysis of CSD indicators

In the stochastic time series analysed here, we first visually identified shifts between E to E/M state and M to E state. Then we took time series segments (the regions marked with boxes in Figs. 2 and 3) prior to a critical transition and examined them for the presence of EWS. For stationarity in residuals, we used Gaussian detrending before performing any statistical analysis of the data. The residuals were then used to calculate the EWSs variance, lag-1 autocorrelation and conditional heteroskedasticity. The time series analysis have been performed using the “Early Warning Signals Toolbox” (http://www.early-warning-signals.org/). A concurrent rise in the varaince and/or lag-1 autocorrelation forewarn an upcoming regime shift. The indicator conditional heteroskedasticity also works similarly (for details see *SI Text, Section 4*).

## Author contributions

S.K.S., H.L., M.K.J., and P.S.D designed research; S.S., S.K.S., M.K.J., and P.S.D. performed research; S.S., S.K.S., H.L., M.K.J., and P.S.D. analyzed data; and S.K.S., M.K.J., and P.S.D. wrote the paper.

## Acknowledgements

S.S. acknowledges the financial support from DST, India under the scheme DST-Inspire (IF160459). S.K.S. is supported by SERB, Department of Science and Technology, Government of India (ECR/2018/000514). M.K.J. is also supported by Ramanujan Fellowship awarded by SERB, Department of Science and Technology, Government of India (SB/S2/RJN-049/2018).

