## Supplementary Information for "Anticipating critical transitions in epithelial-hybrid-mesenchymal cell-fate determination"

#### This PDF file includes:

Supplementary text
Figs. S1 to S4
Tables S1 to S4
References for SI reference citations

### Supporting Information Text

### 1. The chimeric circuit

The core regulatory circuit is composed with four components–two families of E-box binding transcription factors (TFs): SNAIL and ZEB, and two families of microRNAs (miRs): miR-34 and miR-200. In Figure 1A (main text) we have presented a schematic diagram of the EMT core regulatory network. It consists of two highly interconnected modules – miR-34/SNAIL and miR-200/ZEB mutually inhibitory microRNA-TF chimeric feedback loop circuit. The cell-fate determination in terms of epithelial vs. mesenchymal phenotype is based on relative concentration levels of microRNAs and TFs. In general, high levels of miR-34 and miR-200 correspond to the epithelial phenotype, and high levels of SNAIL and ZEB correspond to the mesenchymal phenotype. Both TFs repress epithelial-specific gene expression, including E-cadherin (CDH1), the hallmark of the epithelial phenotype, and can also promote the expression of mesenchymal markers such as N-cadherin and vimentin. This network unit is also known to play an important role as "a motor of cellular plasticity," as it is coupled to many key cellular features, including stemness (tumor-initiation potential), proliferation, apoptosis, resistance to therapies, and cell-cell communication (1). Our previous simulations have shown that miR-34/SNAIL feedback loop, by itself, behaves as a noise-buffering integrator, while miR-200/ZEB loop can exhibit tristability, hence enabling the co-existence of three phenotypes - epithelial, mesenchymal and hybrid epithelial/mesenchymal. This tristability is maintained upon coupling of miR-200/ZEB with miR-34/SNAIL; thus, for simplicity, we consider miR-200/ZEB loop driven by SNAIL for our calculations here (2).

### 2. The deterministic Model

In this model (Eq. 2 in the main text) we consider the monostable miR-34/SNAIL acting as a noise-buffering integrator, and hence SNAIL is considered as an external signal to miR-200/ZEB loop. This external signal can be modelled by the excitatory/inhibitory Hill functions. Therefore, the above miR ($\mu$) based chimeric tristable miR-200/ZEB circuit is modelled by three following coupled deterministic equations with miR ($\mu_{200}$) targeting the mRNA ($m_Z$) of a transcribed protein $Z$:

$$\frac{d\mu_{200}}{dt} = g_{\mu_{200}} H^s(Z, \lambda_{Z,\mu_{200}}) H^s(S, \lambda_{S,\mu_{200}}) - Y_{\mu_{200}} - k_{\mu_{200}}\mu_{200}, \quad [1a]$$

$$\frac{dm_Z}{dt} = g_{m_Z} H^s(Z, \lambda_{Z,m_Z}) H^s(S, \lambda_{S,m_Z}) - Y_{m_Z} - k_{m_Z} m_Z, \quad [1b]$$

$$\frac{dZ}{dt} = L - k_Z Z, \quad [1c]$$

where $g_{\mu_{200}}$ and $g_{m_Z}$ are the synthesis rates of $\mu_{200}$ and $m_Z$ respectively, $k_{\mu_{200}}, k_{m_Z}, k_Z$ are the degradation rates of $\mu_{200}$, $m_Z$, $Z$, respectively and $g_Z$ is the translation rate of protein $Z$ for each mRNA in the absence of miR, $H^S$ is the shifted Hill function, defined as $H^S(Z, \lambda) = H^-(Z) + \lambda H^+(Z)$, $H^-(Z) = 1/(1 + (Z/Z_0)^{n_Z})$, $H^+(Z) = 1 - H^-(Z)$ and $\lambda$ is the fold change from the basal synthesis rate due to protein $Z$. The $\mu_{200}$ dependent miR-mRNA couplings for active miR ($Y_{\mu_{200}}$) and mRNA ($Y_{m_Z}$) degradations, and effective protein translation ($L$) are defined as:

$$Y_{m_Z} = m_Z \sum_i B(n,i)\sigma_{im_Z} \frac{\left(\frac{\mu_{200}^0 r_\mu^+}{r_\mu^-}\right)^i}{\left(1 + \frac{\mu_{200}^0 r_\mu^+}{r_\mu^-}\right)^n} \quad [2]$$

$$Y_{\mu_{200}} = m_Z \sum_i B(n,i)\sigma_{i\mu_{200}} \frac{\left(\frac{\mu_{200}^0 r_\mu^+}{r_\mu^-}\right)^i}{\left(1 + \frac{\mu_{200}^0 r_\mu^+}{r_\mu^-}\right)^n} \quad [3]$$

$$L \simeq g_Z m_Z \sum_i B(n,i)\beta_i \frac{\left(\frac{\mu_{200}^0 r_\mu^+}{r_\mu^-}\right)^i}{\left(1 + \frac{\mu_{200}^0 r_\mu^+}{r_\mu^-}\right)^n}, \quad [4]$$

where $B$ is the binomial coefficient. The details of the model can be found elsewhere (2). We performed the bifurcation analysis of this circuit for a fixed level of SNAIL, $S$. The parameters used for this analysis are given in Table S1 and Table S2.

The individual translation rates and active degradation rates used for the calculation are given in Table S3.

### 3. The stochastic Model

It is well established that the fluctuations in the abundance of molecules in living cells can affect its growth. Therefore, it is important to incorporate such fluctuations in models. To introduce the inherent fluctuations into the model (Eq. (1) in SI), we developed a stochastic version of the EMT model based on ZEB-miR200 mutual repression. In this model, we consider a mixture of nine chemical species: miR ($\mu_{200}$), mRNA with the number of bound miR200, $m_i$, with $i$ ranging from 0 to 6, and the protein ZEB, labeled by the symbol, $Z$. We further consider these nine species are changing via ten diffent chemical reactions, those are presented in the Table S4. Geometrically, the change in numbers of each of the species is the random walk of the molecules on a multidimensional state space. In specific, we developed a definite form of the master equation for grand probability function $p$ through an essential modifications of the above Eq. (1). The obtained master equation were used to derive the Fokker-Planck equation.

In general the grand probability distribution function depends on nine different state numbers. However, one can simplify it by considering that the extension and de-extension steps are sufficiently fast so as to ensure equilibrium between the different bound states of the mRNA. We define macrostate by the triplet, $\begin{bmatrix}\mu^0_{200}\\ m_Z\\ Z\end{bmatrix}$, where $\mu^0_{200}$ is the number of free miR molecules that exist in the particular microstate where all the mRNA molecules are completely unbound, i.e $m_0 = m_z$ and $m_i = 0$ for all $i \neq 0$. We called this triplet as the state vector throughout our discussion. Transitions are allowed only to neighboring states. The change in state vectors in effect of molecular events and the transition probabilities along with all the molecular events those happen for this circuit are described in Table S4.

We label a microstate as $\{m_i\}$, where $\sum_{i=0}^n m_i = m_Z$ and in that state $\mu_{200} = \mu^0_{200} - \sum_{i=0}^n im_i$. These states are connected to each other by the rapid complex formation and complex dissociation reactions. To obtain an equation for the time evolution of macrostate probability, $p(\mu^0_{200}, m_Z, Z)$, we consider two aspects of the problem: the relative probabilities of being in the various microstates for a given macrostate; and the transitions between the macrostates as defines as the average of the transitions between the corresponding microstates. We first deal with the relative transition probabilities among various microstates by insisting on detailed balance for the change of one bound miR from one mRNA molecule. To demonstrate it, let us consider a transition from $m_0 = m_Z$ to the state $m_0 = m_Z - 1$, $m_1 = 1$. It is clear that

$$\hat{p}(m_0 = m_Z - 1, m_1 = 1) = m_Z n \frac{\mu^0_{200} r^+_\mu}{r^-_\mu} \hat{p}(m_0 = m_Z) \quad [5]$$

where $\hat{p}$ denotes microstate probability. The factors $m_Z$ and $n$ come from chosing one mRNA, and selecting one of the sites of mRNA for the binding of one miR. Going further, we can derive

$$\begin{aligned}\hat{p}(m_0 = m_Z - 2, m_1 = 2) &= \frac{1}{2}(m_Z - 1)n(\mu^0_{200} - 1)\frac{r^+_\mu}{r^-_\mu}\hat{p}(m_0 = m_Z - 1, m_1 = 1)\\ &= \frac{m_Z(m_Z-1)}{2}\mu^0_{200}(\mu^0_{200} - 1)\left(\frac{r^+_\mu}{r^-_\mu}\right)^2 \hat{p}(m_0 = m_Z)\ . \end{aligned} \quad [6]$$

Similarly,

$$\begin{aligned}\hat{p}(m_0 = m_Z - 1, m_2 = 1) &= \frac{1}{2}(n-1)(\mu^0_{200} - 1)\frac{r^+_\mu}{r^-_\mu}\hat{p}(m_0 = m_Z - 1, m_1 = 1)\\ &= m_Z\frac{n(n-1)}{2}\mu^0_{200}(\mu^0_{200} - 1)\left(\frac{r^+_\mu}{r^-_\mu}\right)^2 \hat{p}(m_0 = m_Z)\ . \end{aligned} \quad [7]$$

Therefore, the general formulla takes the following form

$$\hat{p}(\{m_i\}) = \left(\frac{r^+_\mu}{r^-_\mu}\right)^{\sum im_i} \mu^0_{200}(\mu^0_{200} - 1)(\mu^0_{200} - 2)\ldots(\mu^0_{200} - \sum im_i + 1)\frac{m_Z!}{\prod_i m_i!}\prod_{i=1}^n B(n,i)^{m_i}\hat{p}(m_0 = m_Z), \quad [8]$$

where $B(n,i)$ is the binomial coefficient. To get this in terms of the overall probability of the macrostate, we just need to chose, $\hat{p}(m_0 = m_Z)$, by the normalization condition

$$\sum_{\{m_i\}} \hat{p}(\{m_i\}) = p(\mu^0_{200}, m_Z, Z)\ . \quad [9]$$

For convenience we refer to the normalized coefficients $\hat{p}(\{m_i\})/p(\mu^0_{200}, m_Z, Z)$ with a symbol $\alpha(\{m_i\})$. Just as a check of this formula, note that if we have one RNA molecule $M = 1$, then the configuration is just decided on by which $j$ state is occupied; if we then neglect the loss of microRNA, we recover the standard binding formula for the normalized probabilities

$$\hat{p}(j) = \alpha(\{m_i\}) = B(n,j)\frac{\left(\frac{r^+_\mu \mu^0_{200}}{r^-_\mu}\right)^j}{\left(1 + \frac{r^+_\mu \mu^0_{200}}{r^-_\mu}\right)^n} \quad [10]$$

Now that we know the relative probabilities of being in each microstate, we can proceed to determine the macrostate transition probabilities. Reaction #1. is simple in that its rate is not dependent on a specific microstate and all transitions take place between the same two macrostates. In other words, we just have a contribution to the master equation.

$$\frac{dp}{dt} = g_{\mu_{200}}(Z)\left(p(\mu^0_{200} - 1, m_Z, Z) - p(\mu^0_{200}, m_Z, Z)\right)\ . \quad [11]$$

This then adds terms to the Fokker-Planck equation as

$$\frac{dp}{dt} = g_{\mu_{200}}(Z)\left(-\frac{\partial p(\mu_{200}^0, m_Z, Z)}{\partial \mu_{200}^0} + 1/2\frac{\partial^2 p(\mu_{200}^0, m_Z, Z)}{\partial {\mu_{200}^0}^2}\right), \quad [12]$$

where $g_{\mu_{200}}(Z)$ is the assumed miRNA transcription rate as a function of the ZEB protein.

Adding a messenger RNA Reaction #2. is also straightforward and just gives

$$\frac{dp}{dt} = g_{m_Z}\left(p(\mu_{200}^0, m_Z - 1, Z) - p(\mu_{200}^0, m_Z, Z)\right) . \quad [13]$$

This then adds terms to the Fokker-Planck equation as

$$\frac{dp}{dt} = g_{m_Z}\left(-\frac{\partial p(\mu_{200}^0, m_Z, Z)}{\partial m_Z} + 1/2\frac{\partial^2 p(\mu_{200}^0, m_Z, Z)}{\partial {m_Z}^2}\right), \quad [14]$$

where $g_{m_Z}$ is the assumed constant mRNA production rate, which is further modulated by ZEB protein.

Reaction #3., Baseline destruction of messenger RNA which leaves all the microRNA intact is also straightforward

$$\frac{dp}{dt} = k_{m_Z}\left((m_Z + 1)p(\mu_{200}^0, m_Z + 1, Z) - m_Z\ p(\mu_{200}^0, m_Z, Z)\right) . \quad [15]$$

This then adds terms to the Fokker-Planck equation as

$$\frac{dp}{dt} = k_{m_Z}\left(\frac{\partial\ m_Z p(\mu_{200}^0, m_Z, Z)}{\partial m_Z} + 1/2\frac{\partial^2\ m_Z p(\mu_{200}^0, m_Z, Z)}{\partial {m_Z}^2}\right) \quad [16]$$

where $k_{m_Z}$ is the assumed constant mRNA decay rate.

The protein decay term #4. looks just like the mRNA decay except that the variable which is changing to $Z$. Hence

$$\frac{dp}{dt} = k_Z\left((Z + 1)p(\mu_{200}^0, m_Z, Z + 1) - Zp(\mu_{200}^0, m_Z, Z)\right) \quad [17]$$

This then adds terms to the Fokker-Planck equation as

$$\frac{dp}{dt} = k_Z\left(\frac{\partial\ Zp(\mu_{200}^0, m_Z, Z)}{\partial Z} + 1/2\frac{\partial^2\ Zp(\mu_{200}^0, m_Z, Z)}{\partial Z^2}\right) \quad [18]$$

where $k_Z$ is the assumed constant mRNA decay rate.

Reaction #5. is a decay of a miRNA molecule at a rate which is independent of whether or not it is bound to a messenger RNA molecule. Hence since all microstates belonging to a given macrostate has the same total miRNA ($= \mu_{200}^0$), we get the simple answer

$$\frac{dp}{dt} = k_{\mu_{200}}\left((\mu_{200}^0 + 1)p(\mu_{200}^0 + 1, m_Z, Z) - \mu_{200}^0 p(\mu_{200}^0, m_Z, Z)\right) . \quad [19]$$

This then adds terms to the Fokker-Planck equation as

$$\frac{dp}{dt} = k_{\mu_{200}}\left(\frac{\partial\ \mu_{200}^0 p(\mu_{200}^0, m_Z, Z)}{\partial \mu_{200}^0} + 1/2\frac{\partial^2 \mu_{200}^0 p(\mu_{200}^0, m_Z, Z)}{\partial {\mu_{200}^0}^2}\right), \quad [20]$$

where $k_{\mu_{200}}$ is the assumed constant miRNA decay rate.

The more complicated reactions are the ones for which the rate depend on the specific microstate. Let us start with reaction #6. protein production and assume that a mRNA molecule with $i$ bound microRNA's will have a translation rate of $\beta_i g_Z$. The full theory here will give rise to a term in the master equation

$$\frac{dp}{dt} = L(\mu_{200}^0, m_Z)\left(p(\mu_{200}^0, m_Z, Z - 1) - p(\mu_{200}^0, m_Z, Z)\right), \quad [21]$$

where

$$L = g_Z \sum_{\{m_i\}}\left(\sum(\beta_j m_j)\right)\alpha(\{m_i\}) . \quad [22]$$

If we want to use a Fokker-Planck equation because the fluctuations are small, we can approximate the above expression for $\alpha$ by replacing

$$\mu_{200}^0(\mu_{200}^0 - 1)(\mu_{200}^0 - 2)...(\mu_{200}^0 - \sum_i im_i + 1) \simeq \mu_0^{\sum_i im_i} . \quad [23]$$

Ignoring the reduction of available microRNA means that each mRNA molecule and each binding site on an individual molecule behaves indepedently. With this approximation, one can show that the expression for the rate reduces to the one given in (2), and this term becomes

$$L \;\simeq\; g_Z m_Z \sum_i B(n,i)\beta_i \frac{\left(\frac{\mu_{200}^0 r_\mu^+}{r_\mu^-}\right)^i}{\left(1+\frac{\mu_{200}^0 r_\mu^+}{r_\mu^-}\right)^n}, \qquad [24]$$

where $B$ is the binomial coefficient. The FP contribution is then

$$\frac{dp}{dt} \;=\; L(\mu_{200}^0, m_Z)\left(-\frac{\partial p(\mu_{200}^0, m_Z, Z)}{\partial Z} + 1/2\frac{\partial^2 p(\mu_{200}^0, m_Z, Z)}{\partial Z^2}\right)\;. \qquad [25]$$

We next go to the enhanced decay of messenger RNA and microRNA if they are bound. We will use the same approximation as given in Eq. (23) above. We can assume different processes that destroy mRNA molecules at rate $\sigma_{im_Z}$ without eliminating any microRNA in addition to miRNA molecules at rate $\sigma_{i\mu_{200}}$ without changing any mRNA molecules. In the Fokker-Planck limit these give the same first derivative terms if $\sigma_{im_Z} = \sigma_{i\mu_{200}}$ but different second derivative terms. The full expression for this term is the complicated contribution

$$\frac{dp}{dt} \;=\; \sum_{j=0}^{n}\left(\Lambda_{jm_Z}(\mu_{200}^0, m_Z+1)p(\mu_{200}^0, m_Z+1, Z) - \Lambda_{jm_Z}(\mu_{200}^0, m_Z)p(\mu_{200}^0, m_Z, Z)\right) \qquad [26]$$

in additon to

$$\frac{dp}{dt} \;=\; \sum_{j=0}^{n}\left(\Lambda_{j\mu_{200}}(\mu_{200}^0+j, m_Z)p(\mu_{200}^0+j, m_Z, Z) - \Lambda_{j\mu_{200}}(\mu_{200}^0, m_Z)p(\mu_{200}^0, m_Z, Z)\right) \qquad [27]$$

with the two different rate factors

$$\Lambda_{jm_Z} \;=\; \sum_{\{m_i\}} \alpha(\{m_i\})\sigma_{jm_Z} m_j \;,\; \Lambda_{j\mu_{200}} \;=\; \sum_{\{m_i\}} \alpha(\{m_i\})\sigma_{j\mu_{200}} m_j \qquad [28]$$

Finally in the Fokker Planck limit and using the approximation we have mentioned, we have the following form

$$\frac{dp}{dt} \;=\; \frac{\partial\, Y_{m_Z}(\mu_{200}^0, m_Z)P(\mu_{200}^0, m_Z, Z)}{\partial m_Z} + 1/2\frac{\partial^2\, Y_{m_Z}(\mu_{200}^0, m_Z)p(\mu_{200}^0, m_Z, Z)}{\partial {m_Z}^2} \qquad [29]$$

$$+\; \frac{\partial\, Y_{\mu_{200}}(\mu_{200}^0, m_Z)p(\mu_{200}^0, m_Z, Z)}{\partial \mu_{200}^0} + 1/2\frac{\partial^2\, Y'_{\mu_{200}}(\mu_{200}^0, m_Z)p(\mu_{200}^0, m_Z, Z)}{\partial {\mu_{200}^0}^2} \qquad [30]$$

with the deterministic expressions found by (2) of

$$Y_{m_Z} \;=\; m_Z \sum_i B(n,i)\sigma_{im_Z} \frac{\left(\frac{\mu_{200}^0 r_\mu^+}{r_\mu^-}\right)^i}{\left(1+\frac{\mu_{200}^0 r_\mu^+}{r_\mu^-}\right)^n} \qquad [31]$$

$$Y_{\mu_{200}} \;=\; m_Z \sum_i B(n,i)\sigma_{i\mu_{200}} \frac{\left(\frac{\mu_{200}^0 r_\mu^+}{r_\mu^-}\right)^i}{\left(1+\frac{\mu_{200}^0 r_\mu^+}{r_\mu^-}\right)^n} \qquad [32]$$

$$Y'_{\mu_{200}} \;=\; m_Z \sum_i iB(n,i)\sigma_{i\mu_{200}} \frac{\left(\frac{\mu_{200}^0 r_\mu^+}{r_\mu^-}\right)^i}{\left(1+\frac{\mu_{200}^0 r_\mu^+}{r_\mu^-}\right)^n} \qquad [33]$$

The relative probabilities of being in each microstate, we can proceed to determine the macrostate transition probabilities. Following the molecular events as described in Table S4, we can write the master equation for the $p(\mu_{200}^0, m_Z, Z)$

$$\begin{aligned}
\frac{\partial p}{\partial t} &= g_{\mu_{200}}(Z)\left(p(\mu_{200}^0-1, m_Z, Z) - p(\mu_{200}^0, m_Z, Z)\right) + g_{m_Z}\left(p(\mu_{200}^0, m_Z-1, Z) - p(\mu_{200}^0, m_Z, Z)\right)\\
&+ k_{m_Z}\left((m_Z+1)p(\mu_{200}^0, m_Z+1, Z) - m_Z\; p(\mu_{200}^0, m_Z, Z, \mu_0)\right) + k_Z\left((Z+1)p(\mu_{200}^0, m_Z, Z+1) - Zp(\mu_{200}^0, m_Z, Z)\right)\\
&+ k_{\mu_{200}}\left((\mu_{200}^0+1)p(\mu_{200}^0+1, m_Z, Z) - \mu_0 p(\mu_{200}^0, m_Z, Z)\right) + L(\mu_{200}^0, m_Z)\left(p(\mu_{200}^0, m_Z, Z-1) - p(\mu_{200}^0, m_Z, Z)\right)\\
&+ \sum_{j=0}^{n}\left(\Lambda_{jm_Z}(\mu_{200}^0, m_Z+1)p(\mu_{200}^0, m_Z+1, Z) - \Lambda_{jm_Z}(\mu_{200}^0, m_Z)p(\mu_{200}^0, m_Z, Z)\right)\\
&+ \sum_{j=0}^{n}\left(\Lambda_{j\mu_{200}}(\mu_{200}^0+j, m_Z)p(\mu_{200}^0+j, m_Z, Z) - \Lambda_{j\mu_{200}}(\mu_{200}^0, m_Z)p(\mu_{200}^0, m_Z, Z)\right)\;.
\end{aligned} \qquad [34]$$

The corresponding Fokker-Planck equation takes the following form for this circuit.

$$
\begin{aligned}
\frac{\partial p}{\partial t} &= g_{\mu_{200}}(Z)\left(-\frac{\partial p(\mu_{200}^0, m_Z, Z)}{\partial \mu_{200}^0} + 1/2\frac{\partial^2 p(\mu_{200}^0, m_Z, Z)}{\partial {\mu_{200}^0}^2}\right) + g_{m_Z}\left(-\frac{\partial p(\mu_{200}^0, m_Z, Z)}{\partial m_Z} + 1/2\frac{\partial^2 p(\mu_{200}^0, m_Z, Z)}{\partial {m_Z}^2}\right) \\
&+ k_{m_Z}\left(\frac{\partial\ m_Z p(\mu_{200}^0, m_Z, Z)}{\partial m_Z} + 1/2\frac{\partial^2\ m_Z p(\mu_{200}^0, m_Z, Z)}{\partial {m_Z}^2}\right) + k_Z\left(\frac{\partial\ Z p(\mu_{200}^0, m_Z, Z)}{\partial Z} + 1/2\frac{\partial^2\ Z p(\mu_{200}^0, m_Z, Z)}{\partial Z^2}\right) \\
&+ k_{\mu_{200}}\left(\frac{\partial\ \mu_{200}^0 p(\mu_{200}^0, m_Z, Z)}{\partial \mu_{200}^0} + 1/2\frac{\partial^2 \mu_{200}^0 p(\mu_{200}^0, m_Z, Z)}{\partial {\mu_{200}^0}^2}\right) + L(\mu_{200}^0, m_Z)\left(-\frac{\partial p(\mu_{200}^0, m_Z, Z)}{\partial Z} + 1/2\frac{\partial^2 p(\mu_{200}^0, m_Z, Z)}{\partial Z^2}\right) \\
&+ \frac{\partial\ Y_{m_Z}(\mu_{200}^0, m_Z) P(\mu_{200}^0, m_Z, Z)}{\partial m_Z} + 1/2\frac{\partial^2\ Y_{m_Z}(\mu_{200}^0, m_Z) p(\mu_{200}^0, m_Z, Z)}{\partial {m_Z}^2} \\
&+ \frac{\partial\ Y_{\mu_{200}}(\mu_{200}^0, m_Z) p(\mu_{200}^0, m_Z, Z)}{\partial \mu_{200}^0} + 1/2\frac{\partial^2\ Y'_{\mu_{200}}(\mu_{200}^0, m_Z) p(\mu_{200}^0, m_Z, Z)}{\partial {\mu_{200}^0}^2}. \qquad [35]
\end{aligned}
$$

**A. Monte Carlo simulation .** To obtain the influence of intrinsic noise on dynamical properties of the system, one needs to solve the above master equation (Eq. (34)) by analytical approach. The complexity of the master equation associated with a dynamical system does not allow to solve this equation analytically for most of the cases. However, the master equation can be exactly solved by numerical approach by means of Monte Carlo simulation. We used Gillespie algorithm to perform our simulation. Monte Carlo simulation provides individual realizations of Markov process. It has wide range of applications in the fields like systems biology, ecology, chemical reaction dynamics, etc.

In Table S4, we have stated all the individual associated processes those exhibit in our system. The change in state vector, $\begin{bmatrix}\mu_{200}^0 \\ m_Z \\ Z\end{bmatrix}$ and the propensity function, $a_n$ are also mentioned in the Table S4. Using this elementary events with the parameters, we have performed Monte Carlo simulation and updated the corresponding miRNA ($\mu_{200}^0$), mRNA ($m_Z$) and protein ($Z$). For this simulation we have used direct Gillespie algorithm that first calculate the propensity function then draw two random numbers ($r_1$ and $r_2$) from an uniform distribution in the interval $(0, 1)$. The first random number, $r_1$ is used to stochastic update of time $(t + \tau)$ during a next reaction occur, where

$$\tau = \frac{1}{a_0}\ln(\frac{1}{r_1})$$

with $a_0 = \sum_{i=1}^{n} a_j$. The index $n$ of the occurring process is given by the smallest integer satisfying:

$$\sum_{i=1}^{n-1} a_j < r_2 a_0 \leq \sum_{i=1}^{n} a_j.$$

The system states are updated $X(t+\tau) = X(t)+\Delta X_{\nu_n}$, then the simulation proceeds to the next occurring time. $\Delta X_{\nu_n}$ represents change in species vector in the stochastic time interval $\tau$.

### 4. Early warning indicators

**A. Variance and Autocorrelation.** We consider simulated stochastic time series associated with regime shifts to calculate the statistical indicators. Gaussian detrending with bandwidth 85, 80 and 80 have been used to get stationary residuals to the time series corresponding to: E to hybrid-E/M transition, M to E reverse transition, and hybrid-E/M to M state transition, respectively. To anticipate critical transition, we first calculate variance and autocorrelation at lag-1 of the residual time series in a moving window. Autocorrelation at lag-1 can be calculated from the autocorrelation function (ACF), defined as:

$$\rho_t = \frac{E\left(m'(t), m'(t+1)\right)}{var(m(t))},$$

where $m'(t) = m(t) - E(m(t))$, and $m(t)$ represents mRNA concentration at a time instant $t$ and $var(m(t))$ is the variance of mRNA. Variance is calculated as:

$$var(m(t)) = \frac{1}{n}\sum\left(m(t) - E(m(t))\right)^2,$$

where $E\left(m(t)\right)$ and $n$ represent the mean value and total number of observations in the data set, respectively.

**B. Sensitivity Analysis.** The contour plots are representative to the sensitivity analysis for lag-1 autocorrelation and standard deviation. It represents that for different combination of window size and bandwidth, how trend statistic, Kendall $\tau$ is distributed and where it will give more sensitive results. Kendall $\tau$ is the correlation between the two parameters bandwidth and window size. This analysis gives an idea about the robustness of choices of window size and bandwidth. Window sizes ranging from 25% to 75% of the time series length with increments of 15 points has been used for evaluating lag-1 autocorrelation and for bandwidth, ranges from 5 to 100 with increments of 10. An increasing trend for both the lag-1 autocorrelation and variance is observed for our analysis.

**C. Conditional Heteroskedasticity.** To reduce the chance of false positive signals form the time series, we have also evaluated the conditional heteroskedasticity which is one of the robust indicators of critical slowing down (3). To check whether the time series is heteroskedastic or not, we have plotted squared residuals at a time step $t$ vs. the time step $t+1$ and did cumulative significant test of the time series prior to a critical transition. While calculating conditional heteroskedasticity, we took window size of 10% of the data and then apply Gaussian detrending, and calculate residual square. For the cumulative significant test, the procedure is as follows:

First we take the time series prior to the transition. Lag the data one time step and fit an AR(1) model using least squares regression. Calculate squared residuals from the AR(1) model. Then calculate The Lagrange Multiplier test statistic. Finally to check whether the test statistic is giving the positive result or not, compare this to a chi-square distribution with one degree of freedom (3).

### 5. Transition from hybrid-E/M state to M state and the corresponding early warning signals

Here we investigate the EWS to forewarn a sudden critical transition from hybrid-E/M state to M state with increasing level of SNAIL. For the analysis we consider a time series segment before the transition from hybrid-E/M state to M state (Fig. S1(A)). To remove the non stationarity in the data we first fit a Gaussian kernal smoothing function with a bandwidth of 80 and then subtract the curve from the time series segment. The remaining residuals (Fig. S1(C)) time series has been used for the EWS analysis. We calculate the variance and AR(1) with a rolling window size of length 70% the length of the residual time series segment, and found both the variance and AR(1) are increasing. This shows EWS also work well for this transition. Also we did the sensitivity analysis for choosing the bandwidth and window size. For sensitivity analysis the rolling window size was varied from 25% to 75% of the data length in increments of 15 points, together with variations in the filtering bandwidth ranging from 5 to 100 in increments of 10 points. This sensitivity analysis is quantified using the non-parametric Kendall's $\tau$ rank correlation coefficient. A positive Kendall's $\tau$ determines increasing trend in the EWS prior to a critical transition. To maximise the estimated trends for the EWS, we have used the sensitivity plot to select a particular filtering bandwidth and window size (see Fig. S1(F) for variance and Fig. S1(H) for AR(1)).

### 6. Box plot

For a particular filtering bandwidth and a combination of rolling window size, how the time series prior to the transition is sensitive is measured by the Kendall's $\tau$ rank correlation coefficient. The box plot "org" shows the distribution of trends for the time series. To test the significance, we generate 100 surrogate time series using ‘ARMA‘ model then calculate trend statistic Kendall's $\tau$ and show the distribution of trends represented by "sg". Trends distribution for "org" & "sg" and mean differences shows how much data based trends differ from the surrogate based trends. Fig. S2(A) and Fig. S2(B) depict the sensitivity and the significance for the indicators AR(1) and variance, respectively. To generate this surrogate time series from the original time series we have used 'autoregressive moving average' method, i.e. here first we fit a ARMA($p, q$) curve to the time series. This model contains the AR($p$) and MA($q$) models, which is given by

$$X_t = c + \epsilon_t + \sum_{i=1}^{p} \phi_i X_{t-1} + \sum_{i=1}^{q} \theta_i \epsilon_{t-i},$$

where $\phi_1, \phi_2, ..., \phi_p$ and $\theta_1, \theta_2, ..., \theta_q$ are parameters, c is a constant, $\epsilon_t$ is a white noise and $\epsilon_t, \epsilon_{t-1}, ..., \epsilon_{t-q}$ are again white noise error terms. The error terms are generally assumed to be independent identically distributed and sampled from normal distribution with zero mean.

### 7. Stochastic potential

We calculated stochastic potential numerically by using the trajectory obtained from the Monte Carlo simulations. Each of the simulations were performed for about 20 days period to reach steady state for 20000 different values of initial conditions at three values of SNAIL numbers (i.e. S=188K, 200K, and 213K). Once we obtained the steady state trajectory, we calculated frequency count of visits on the rectengular grid in ZEB mRNA-$\mu$RNA200 plane. If we define the frequency count of visits $\Omega_{\mu_{200}^{ss} m_Z^{ss}}$ at a particular value of steady state numbers of ZEB mRNA ($m_Z^{ss}$) and $\mu$RNA200 ($\mu_{200}^{ss}$), and $\Omega_{total}$ is the total count, then we can define the probability of visit for a specific values of ZEB mRNA and $\mu$RNA200 by $p_{ss}(\Omega_{\mu_{200}^{ss}, m_Z^{ss}}) = \frac{\Omega_{\mu_{200}^{ss} m_Z^{ss}}}{\Omega_{total}}$. The obtained probabilities can be used to calculate the stochastic potential by taking negative logarithm of it, $(F = -\ln(p_{ss}))$. The stability of a state is determined by the lower values of the stochastic potential.

### 8. Basin stability

For a multistable system, basin stability is the measurement of basin volume for different stable states (4). In our system (for a particular value of SNAIL) we have three different stable states, i.e. system have three stable equilibrium points. Measure of basin stability for the given system is as follows: for a particular value of the signal SNAIL (say S=200K), we choose a set of initial conditions, say $I$ which is bounded in $\mathbb{R}^3$. We randomly draw N=10000 initial conditions from $I$ and numerically solve the given deterministic equation for these set of initial conditions. After numerically solving the system and removing enough transients this N uniformly distributed random initial conditions are distributed among this three stable equilibrium point.

Suppose $N_1$ the number of initial conditions that converge to one stable state among these three stable state. Then the basin stability of that particular stable state is estimated as $\frac{N_1}{N}$. In Fig. S3 we have evaluated the basin stability of the chimeric circuit for different values of the signal SNAIL starting with a bistability region (co-existence of E and M) and then moving to tristable region (co-existence of E, E/M, and M) and finally again goes back to a bistable region (co-existence of E/M and M). We observe that the hybrid-E/M state is very stable and becomes the dominated state since its inception, and remains so until it exists. This analysis challenges the long-held assumption about hybrid E/M state(s) being 'metastable', instead suggests that cells can stably attain and maintain these hybrid E/M phenotypes (5).

### 9. Phase diagram with variations in the signal SNAIL and the threshold for ZEB self activation levels

To identify other potential parameters enabling a monostable hybrid-E/M state, we varied SNAIL and the threshold value of ZEB in the shifted Hill function corresponding to its self-activation (Fig. S4). We again observed four saddle-node bifurcation curves, and identified a monostable hybrid-E/M region at relatively lower values of SNAIL and ZEB self-activation threshold. This result suggests that at low SNAIL values enabling an epithelial state, strengthening ZEB self-activation can switch the cells to being hybrid-E/M; however, increasing SNAIL pushes these hybrid-E/M cells to mesenchymal. Overall, a balance between the strengths of mutual inhibition and self-activation in ZEB/miR-200 loop can enable a monostable hybrid-E/M state. Conversely, disrupting this balance can evade transitions to a hybrid-E/M state. In Fig. S4(B)–(E) bifurcation diagrams of mRNA with variations in the level of SNAIL for different values of threshold for ZEB self activation ($Z_{0,m_z}$): (B) 10K, (C) 20K, (D) 30K, and (E) 40K are presented. For $Z_{0,m_z} = 10$K the monostable hybrid-E/M state exists. With an increase in the $Z_{0,m_z}$ the monostability of hybrid-E/M state disappears and E state appears. Further increase in the value of $Z_{0,m_z}$ results in the decrease of existence region of the hybrid-E/M state.

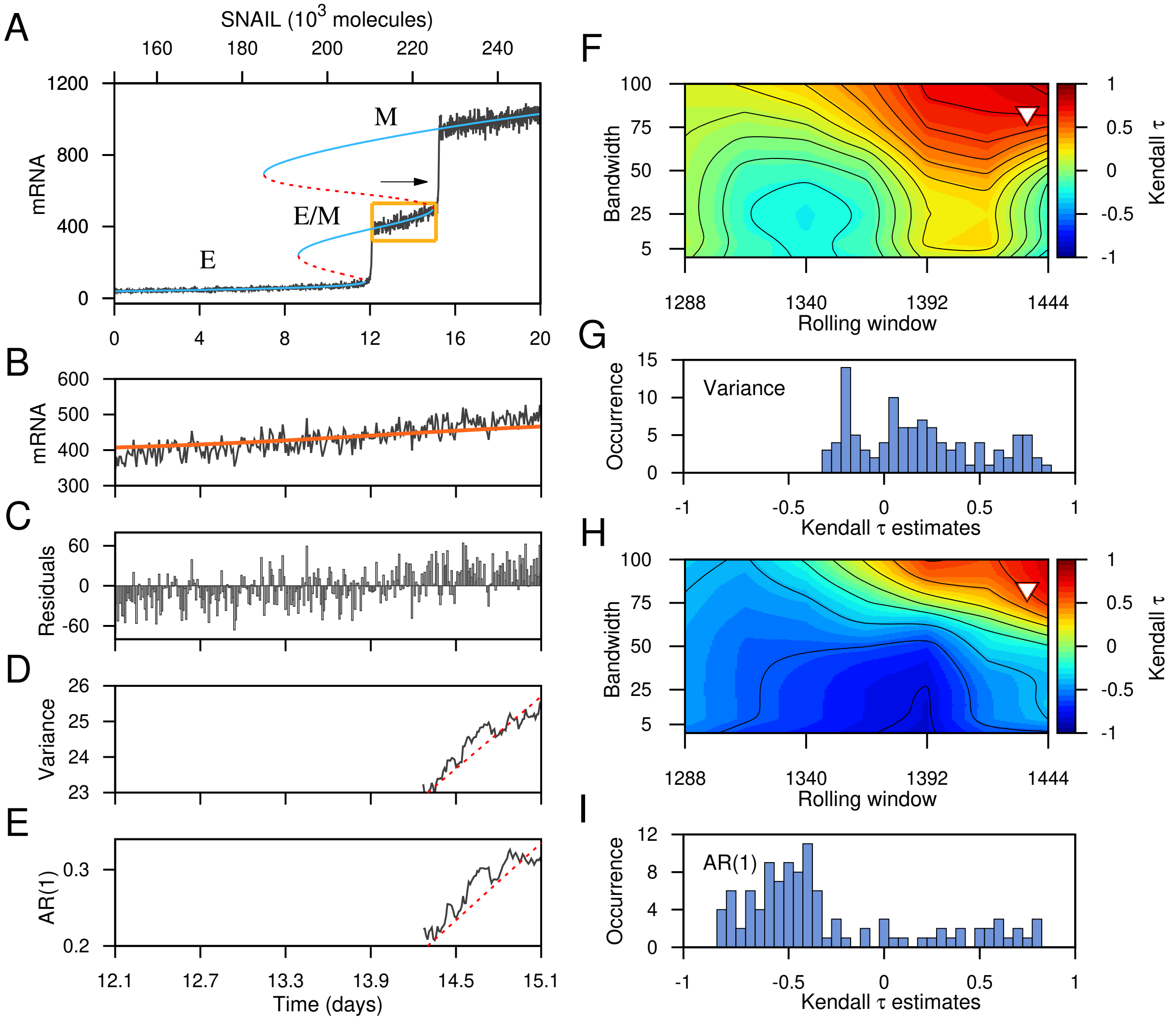


**Fig. S1.** Critical transition between different cell states of the regulatory circuit, driven by forward change of the control parameter SNAIL and indicators of critical slowing down. (A) Transition from E state to Hybrid E/M state and hybrid E/M state to M state. (B) Stochastic time series taken for the analysis as indicated by boxed region in (A). (C) Residual time series after applying Gaussian filtering (red curve in (B) is the trend used for filtering). EWS calculated from the residual time series after using a rolling window size 70% of the data length. (D) variance and (E) AR(1). (F-I) Sensitivity analysis for all combination of filtering bandwidth and rolling window size and used to calculate EWS. Contour plots reveal the effect of rolling window size and filtering bandwidth on the observed trend in the EWS, (F) variance and (H) AR(1), for the filtered data as measured by the Kendall- $\tau$ value. The triangles indicate the rolling window size and bandwidth used to calculate the EWS. Frequency distributions of Kendall-$\tau$ values for (G) variance and (I) AR(1).

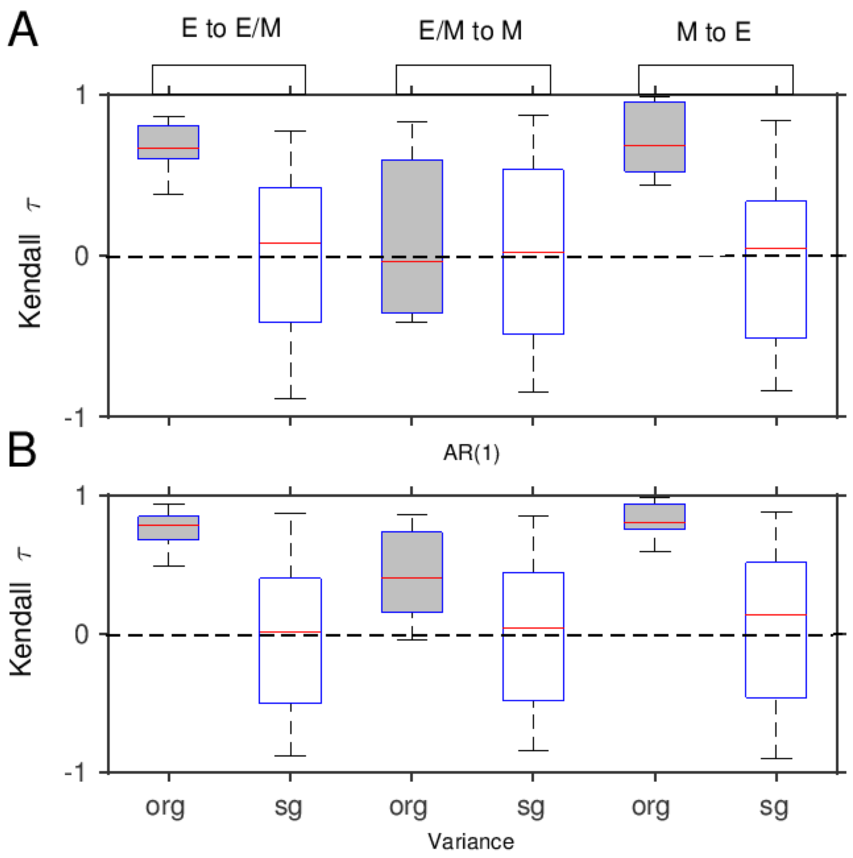


**Fig. S2.** Sensitivity analysis for stochastic time series and a comparison between the trends for all three types of critical transition: E to hybrid E/M, hybrid E/M to M, and M to E. Box plot shows distribution of trend statistic Kendall- $\tau$ across all window sizes for the trends (A) AR(1) and (B) variance. The boxes labeled as 'org' signifies that it's for the original data set before the transition and 'sg' represents distribution of trends based on 100 surrogate time series with the same ARMA structure. Red lines show the median curve for this distribution.

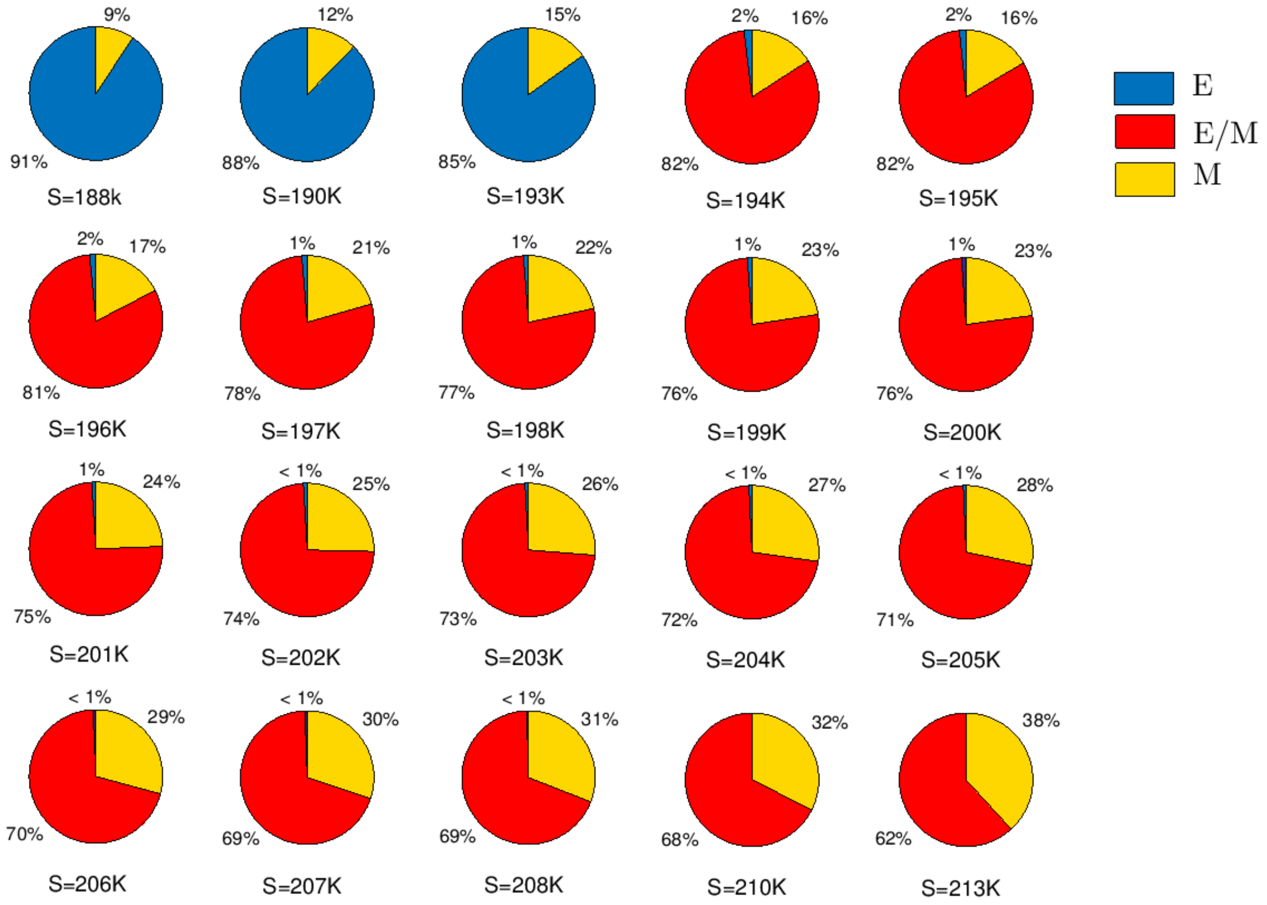


**Fig. S3.** Pie diagrams representing basin stability of the system for different values of the signal SNAIL. The percentage of $10^4$ simulations with random initial conditions reaching to a particular steady state in a bistable/multistable region are presented. Blue, red and yellow regions correspond to the $\%$ of simulations reaching to the E, hybrid-E/M and M states, respectively. Pie diagram shows that in the tristable region, i.e for the SNAIL value S$\approx$194K to S$\approx$208K, hybrid-E/M state is more stable than E or M state. Even beyond the value of 208K, i.e. in the bistable region S$=$210K and S$=$213K hybrid E/M state is more stable.

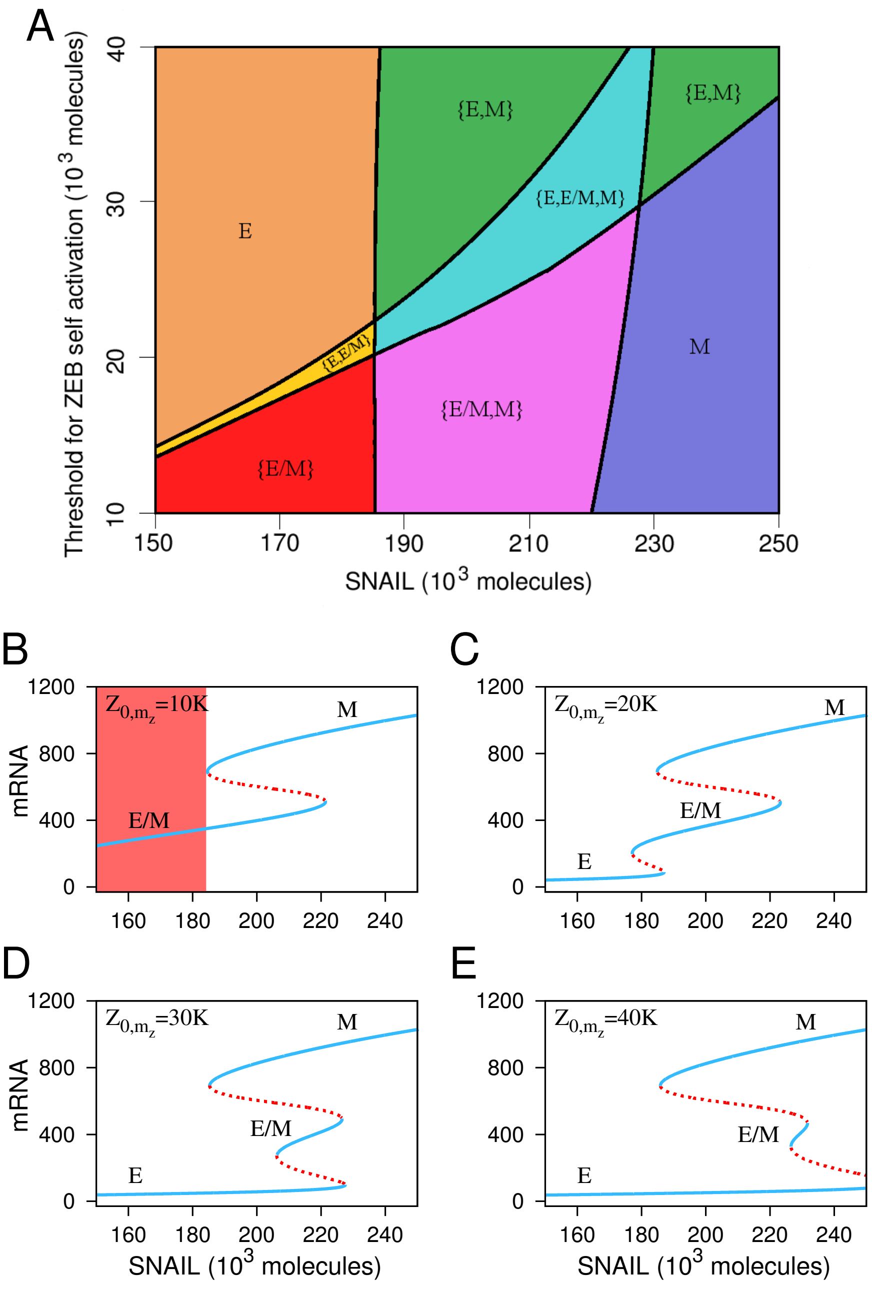


**Fig. S4.** Phase diagram and corresponding bifurcation diagrams of the genetic circuit. (A) The phase diagram of the genetic circuit with variations in levels of SNAIL and threshold for ZEB self activation for miR-200/ZEB= 220K. Each phase corresponds to either any of the monostable state (E, hybrid-E/M and M) or coexisting bistable/tristable states. For example, in E the epithelial state is stable, in {E, hybrid-E/M} both epithelial and hybrid-epithelial-mesenchymal states coexist. (B–E) Bifurcation diagrams of mRNA with variations in the level of SNAIL for different values of threshold for ZEB self activation ($Z_{0,m_z}$): (B) 10K, (C) 20K, (D) 30K, and (E) 40K. For $Z_{0,m_z} = 10$K the monostable hybrid-E/M state exists (B). With an increase in the $Z_{0,m_z}$ the monostability of hybrid-E/M state disappears and E state appears (C). Further increase in the value of $Z_{0,m_z}$ results in the decrease of existence region of the hybrid-E/M state (D, E).

**Table S1. Synthesis and degradation parameters for the miR-200/ZEB circuit**

| Sl. No. | Synthesis rate | Degradation rate |
|---|---|---|
| 1. | $g_{\mu_{200}} = 2.1 \times 10^3 h^{-1}$ | $k_{\mu_{200}} = 0.05 h^{-1}$ |
| 2. | $g_{m_Z} = 11 h^{-1}$ | $k_{m_Z} = 0.5 h^{-1}$ |
| 3. | $g_Z = 0.1 \times 10^3 h^{-1}$ | $k_Z = 0.1 h^{-1}$ |

**Table S2. Hill function parameters for the miR-200/ZEB circuit**

| Sl. No. | Molecules | Coefficients | Fold change |
|---|---|---|---|
| 1. | $Z_{0,\mu_{200}} = 220 \times 10^3$ | $n_{Z,\mu_{200}} = 3$ | $\lambda_{Z,\mu_{200}} = 0.1$ |
| 2. | $Z_{0,m_Z} = 25 \times 10^3$ | $n_{S,\mu_{200}} = 2$ | $\lambda_{S,\mu_{200}} = 0.1$ |
| 3. | $S_{0,\mu_{200}} = 180 \times 10^3$ | $n_{Z,m_Z} = 2$ | $\lambda_{Z,m_Z} = 7.5$ |
| 4. | $S_{0,m_Z} = 180 \times 10^3$ | $n_{S,m_Z} = 2$ | $\lambda_{S,m_Z} = 10$ |

**Table S3. Site dependent parameters of mRNA, $n$, $\sigma_{i\mu_{200}}$, $\sigma_{im_Z}$ and $\beta_i$ of the miR-200/ZEB circuit**

| | | | | | | | |
|---|---|---|---|---|---|---|---|
| $n_{\mu_{200}}$ | 0 | 1 | 2 | 3 | 4 | 5 | 6 |
| $\sigma_{i\mu_{200}}(h^{-1})$ | 0 | 0.005 | 0.05 | 0.5 | 0.5 | 0.5 | 0.5 |
| $\sigma_{im_Z}(h^{-1})$ | 0.0 | 0.04 | 0.2 | 1.0 | 1.0 | 1.0 | 1.0 |
| $\beta_i$ | 1.0 | 0.6 | 0.3 | 0.1 | 0.05 | 0.05 | 0.05 |

**Table S4. Ten different reactions for the miR-200/ZEB circuit, change of state vectors, gain and loss probabilities, and their propensity function. The symbols ($+1$) and ($-1$) in the column of state vectors represent birth and death processes of the respective chemical species. Here, $P$ stands for permutation and $p$ stands for the grand probability function.**

| Sl. No. | Molecular Events | Reaction Type | Before Reaction | After Reaction | Gain Probability | Loss Probability | Propensity Function ($a_n$) |
|---|---|---|---|---|---|---|---|
| 1. | $\phi \xrightarrow{g_\mu} \mu$ | Creation of miR | $\begin{bmatrix}\mu_{200}^0-1\\ m_Z\\ Z\end{bmatrix}$ | $\begin{bmatrix}\mu_{200}^0\\ m_Z\\ Z\end{bmatrix}$ | $g_{\mu_{200}}p(\mu_{200}^0-1, m_Z, Z)$ | $g_{\mu_{200}}p(\mu_{200}^0, m_Z, Z)$ | $g_{\mu_{200}}$ |
| 2. | $\phi \xrightarrow{g_m} m_0$ | Creation of mRNA | $\begin{bmatrix}\mu_{200}^0\\ m_Z-1\\ Z\end{bmatrix}$ | $\begin{bmatrix}\mu_{200}^0\\ m_Z\\ Z\end{bmatrix}$ | $g_{m_Z}p(\mu_{200}^0, m_Z-1, Z)$ | $g_{m_Z}p(\mu_{200}^0, m_Z, Z)$ | $g_{m_Z}$ |
| 3. | $m \xrightarrow{k_m} \phi_m$ | Direct (baseline) degradation of mRNA | $\begin{bmatrix}\mu_{200}^0\\ m_Z+1\\ Z\end{bmatrix}$ | $\begin{bmatrix}\mu_{200}^0\\ m_Z\\ Z\end{bmatrix}$ | $k_{m_Z}(m_Z+1)$ $\times p(\mu_{200}^0, m_z+1, Z)$ | $k_{m_Z}m_Z p(\mu_{200}^0, m_Z, Z)$ | $k_{m_Z}m_Z$ |
| 4. | $Z \xrightarrow{k_Z} \phi_Z$ | Protein degradation | $\begin{bmatrix}\mu_{200}^0\\ m_Z\\ Z+1\end{bmatrix}$ | $\begin{bmatrix}\mu_{200}^0\\ m_Z\\ Z\end{bmatrix}$ | $k_Z(Z+1)$ $\times p(\mu_{200}^0, m_Z, Z+1)$ | $Zk_Z p(\mu_{200}^0, m_Z, Z)$ | $Zk_Z$ |
| 5. | $\mu \xrightarrow{k_\mu} \phi_\mu$ | Degradation of free miR | $\begin{bmatrix}\mu_{200}^0+1\\ m_Z\\ Z\end{bmatrix}$ | $\begin{bmatrix}\mu_{200}^0\\ m_Z\\ Z\end{bmatrix}$ | $k_{\mu_{200}}(\mu_{200}^0+1)$ $\times p(\mu_{200}^0+1, m_Z, Z)$ | $k_{\mu_{200}}\mu_{200}^0 p(\mu_{200}^0, m_Z, Z)$ | $k_{\mu_{200}}\mu_{200}^0$ |
| 6. | $m_i \xrightarrow{g_Z} Z + m_i$ | Protein production | $\begin{bmatrix}\mu_{200}^0\\ m_Z\\ Z-1\end{bmatrix}$ | $\begin{bmatrix}\mu_{200}^0\\ m_Z\\ Z\end{bmatrix}$ | $L(\mu_{200}^0, m_Z)$ $\times p(\mu_{200}^0, m_Z, Z-1)$ | $L(\mu_{200}^0, m_Z)p(\mu_{200}^0, m_Z, Z)$ | $L(\mu_{200}^0, m_Z)$ |
| 7. | $m_i \rightarrow \phi_m + i\mu$ | Degradation of mRNA (proteolysis) | $\begin{bmatrix}\mu_{200}^0\\ m_Z+1\\ Z\end{bmatrix}$ | $\begin{bmatrix}\mu_{200}^0\\ m_Z\\ Z\end{bmatrix}$ | $\Lambda_{jm_Z}(\mu_{200}^0, m_Z+1)$ $\times p(\mu_{200}^0, m_Z+1, Z)$ | $\Lambda_{jm_Z}(\mu_{200}^0, m_Z)$ $\times p(\mu_{200}^0, m_Z, Z)$ | $\Lambda_{jm_Z}(\mu_{200}^0, m_Z)$ |
| 8. | $m_i \rightarrow m_0 + i\phi_\mu$ | Degradation of miR (proteolysis) | $\begin{bmatrix}\mu_{200}^0+i\\ m_Z\\ Z\end{bmatrix}$ | $\begin{bmatrix}\mu_{200}^0\\ m_Z\\ Z\end{bmatrix}$ | $\Lambda_{j\mu_{200}}(\mu_{200}^0+i, m_Z)$ $\times p(\mu_{200}^0+i, m_Z, Z)$ | $\Lambda_{j\mu_{200}}(\mu_{200}^0, m_Z)$ $\times p(\mu_{200}^0, m_Z, Z)$ | $\Lambda_{j\mu_{200}}(\mu_{200}^0, m_Z)$ |
| 9. | $\mu + m_{i-1} \xrightarrow{r_\mu^+} m_i$ | Complex extension of miR-mRNA | $\begin{bmatrix}\mu_{200}^0+1\\ m_Z\\ Z\end{bmatrix}$ | $\begin{bmatrix}\mu_{200}^0\\ m_Z\\ Z\end{bmatrix}$ | $m_Z r_\mu^+(\mu_{200}^0+1)$ $\times p(\mu_{200}^0+1, m_Z, Z)$ | $m_Z r_\mu^+ \mu_{200}^0$ $\times p(\mu_{200}^0, m_Z, Z)$ | $m_Z r_\mu^+ \mu_{200}^0$ |
| 10. | $m_i \xrightarrow{r_\mu^-} \mu + m_{i-1}$ | Complex de-extension of miR-mRNA | $\begin{bmatrix}\mu_{200}^0-1\\ m_Z\\ Z\end{bmatrix}$ | $\begin{bmatrix}\mu_{200}^0\\ m_Z\\ Z\end{bmatrix}$ | $m_Z r_\mu^-(\mu_{200}^0-1)$ $\times p(\mu_{200}^0-1, m_Z, Z)$ | $m_Z r_\mu^-(\mu_{200}^0)$ $\times p(\mu_{200}^0, m_Z, Z)$ | $m_Z r_\mu^-(\mu_{200}^0)$ |
